# Synthetic microbial co-cultures for modular bioelectronic sensing in diverse environments

**DOI:** 10.1101/2025.09.24.678173

**Authors:** Siliang Li, Duolong Zhu, Kundan Saha, Biki Bapi Kundu, Sameer Sonkusale, Robert A. Britton, Caroline M. Ajo-Franklin

**Affiliations:** Department of BioSciences, Rice University, Houston, TX, USA; Department of Molecular Virology and Microbiology, Baylor College of Medicine, Houston, TX, USA; Department of Electrical and Computer Engineering, Tufts University, Medford, MA, USA; Sonkusale Research Labs, Tufts University, Medford, MA, USA; PhD Program in Systems, Synthetic, and Physical Biology, Rice University, Houston, TX, USA; Alkek Center for Metagenomics and Microbiome Research, Baylor College of Medicine, Houston, TX, USA; Dan L. Duncan Comprehensive Cancer Center, Baylor College of Medicine, Houston, TX, USA; Department of Chemical and Biomolecular Engineering, Rice University, Houston, TX, USA; Department of Bioengineering, Rice University, Houston, TX, USA; Rice Synthetic Biology Institute, Rice University, Houston, TX, USA

## Abstract

Human disruption of ecosystems poses a significant threat to global health, driving the need for low-cost, low-power, and easily deployable sensors for environmental and health monitoring. Microbial bioelectronic sensors are particularly well-suited as they generate electrical signals and can be integrated into compact electronic devices for field deployment over extended periods. However, current engineering strategies for bioelectronic sensors lack modularity, are limited to a few microbial chassis, and depend on specialized instruments for signal detection. Here, we present the <u>e</u>lectroactive <u>co</u>-culture <u>sen</u>sing <u>s</u>ystem (e^-^COSENS), a plug-and-play platform for bioelectronic sensor development. This system comprises a “sender” bacterium that produces electron mediators in response to analytes and a “receiver” bacterium that utilizes the electron mediators to generate electrical signals via <u>e</u>xtracellular <u>e</u>lectron transfer (EET). By modularly swapping the sender bacterium and its associated genetic sensing elements, we achieved bioelectronic sensing of metals, small molecules, and peptides in distinct environmental, food, and human-relevant settings. Moreover, we designed a centimeter-sized bioelectronic device that enables low-cost, portable signal readout from e^-^COSENS using a household digital multimeter. The e^-^COSENS platform greatly simplifies the bioelectronic sensor design and opens unprecedented potential for bioelectronic sensor applications.

## Introduction

The rapidly expanding global human footprint has caused severe pollution of the environmental and food chains^1^, necessitating continuous monitoring of pollutants and health parameters using low-cost and deployable sensing techniques. Synthetic biology has engineered microbes as whole-cell biosensors to detect chemicals in complex environments over extended periods through analyte-inducible genetic circuits and generate optical^2^, acoustic^3^, or electrical signals^4^. Among these, bioelectronic sensors— which generate electrical signals—are especially well-suited as they rely on low-cost and low-power electronic components for signal measurement^4^ and can detect analytes within minutes of contact even in opaque environments^5^. Bioelectronic sensors have been developed to sense pollutants^5–7^ and pharmacologically relevant molecules^8,9^, and can be integrated into miniature electronic devices for use in urban waterways^5,10–13^, sediments^14,15^, or on human skin^16^, showing promise as on-demand tools for sensing in urban settings.

However, the development of whole-cell biosensors with electrical output lags far behind other biosensor techniques, such as fluorescent biosensors^2^, due to complex engineering strategies and the shortage of suitable microbial chassis^4^. Current approaches typically modulate the on-off activity of <u>e</u>xtracellular <u>e</u>lectron transfer (EET) pathways in native electroactive microbes, such as the mesophilic, freshwater-dwelling *Shewanella oneidensis*^6,7^ or the oxygen-sensitive *Geobacter sulfurreducens*^17^. These approaches restrict the application range of bioelectronic sensors due to the narrow environmental tolerance of native electroactive microbes and limited availability of genetic tools^18^. Synthetic biology allows heterologous expression of multi-component EET pathways in non-native electroactive hosts, such as the model organism *Escherichia coli*^5,19^ or the marine bacterium *Marinobacter atlanticus*^20^. However, these strategies have only succeeded after tedious trial and error yet yield electrical signals far below native systems^21–23^. Furthermore, detecting electrical signals historically requires specialized instruments, such as potentiostats or source meters, which hinders the broader adoption of bioelectronic sensors. Thus, simpler and more efficient engineering and measurement strategies are needed to fully unlock the potential of bioelectronic sensors.

Here, we present the <u>e</u>lectroactive <u>co</u>-culture <u>sen</u>sing <u>s</u>ystem (e^-^COSENS), a plug-and-play platform for creating bioelectronic sensors (**Fig. 1**). This system comprises a “sender” bacterium engineered to produce an electron mediator (1,4-dihydroxy-2-naphthoic acid, DHNA) in response to specific analytes, and a “receiver” bacterium capable of using DHNA as an electron mediator to generate electrical signals through a DHNA-mediated EET pathway^24,25^ (**Fig. 1**). This division of functions enables non-electroactive bacteria to be modularly integrated for versatile bioelectronic sensing. By employing the generally-recognized-as-safe *Lactiplantibacillus plantarum* as a universal receiver^26^, e^-^COSENS demonstrates robust functionality across diverse environmental and human-relevant settings. We also integrate e^-^COSENS into a miniature microbial fuel cell device, which requires only a household digital multimeter for readout, highlighting the possibility of tailoring e^-^COSENS for portable biosensing applications.

**Fig. 1.**
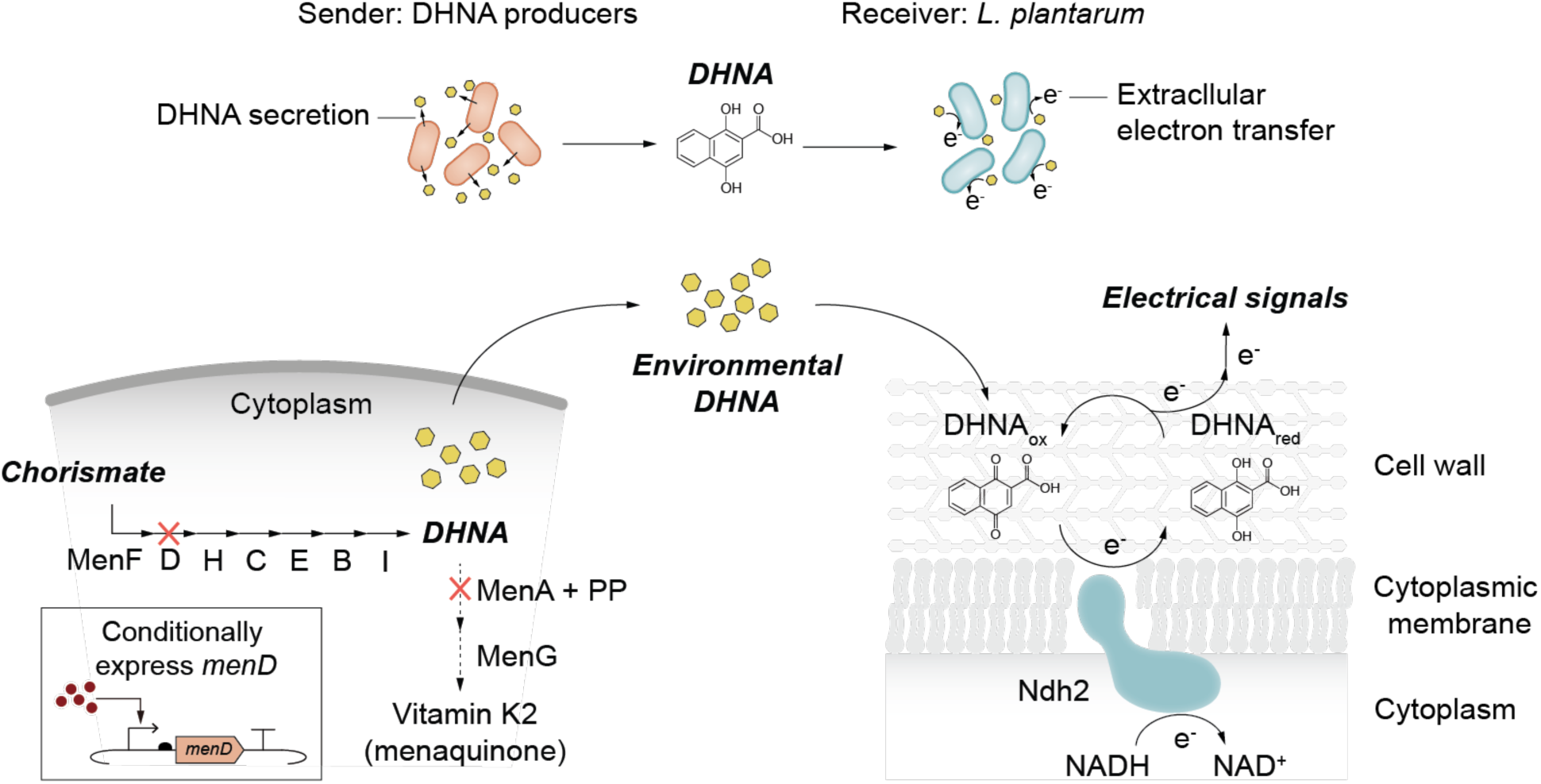
Mechanism of the electroactive co-culture sensing system (e^-^COSENS). The e^-^COSENS system comprises a “sender” and a “receiver” bacterial strain. The sender synthesizes 1,4-dihydroxy-2-naphthoic acid (DHNA, precursor of vitamin K2) from the substrate chorismate through the seven-enzyme Men pathway. The *menD* gene was knocked out from the genome and expressed on plasmids under the control of analyte-inducible promoters to conditionally turn on DHNA biosynthesis for sensing purposes. The *menA* gene was also knocked out to prevent the prenylation of DHNA by prenyl diphosphate (PP). The secreted DHNA activates extracellular electron transfer (EET) in the receiver strain by mediating electron cycling between the membrane-bound type-II NADH:quinone oxidoreductase Ndh2 and the terminal electron acceptors, such as iron(III) oxide or an electrode. This results in the generation of electrical signals. The present system uses *Lactiplantibacillus plantarum* as the receiver strain.

## Results

### “Sender” and “receiver” candidates are widespread

The e^-^COSENS platform is designed based on a quinone-mediated EET pathway possessed by many Gram-positive bacteria^24,25^. The central component is the membrane-bound type-II NADH:quinone oxidoreductase Ndh2, which uses quinones (e.g., DHNA) to shuttle electrons from the cytosolic NADH pool to extracellular electron acceptors, such as iron(III) oxide or electrodes (**Fig. 1**). Studies have shown that certain quinone auxotrophs, such as *L. plantarum*, can perform Ndh2-dependent EET with either exogenous DHNA^8,25,27^ or when incubated with the spent media of a quinone-producing bacterium^28^. This quinone cross-feeding phenomenon inspired us to create sender-receiver co-cultures for EET. As designed, the sender bacterium must be able to produce DHNA and the receiver bacterium must be a DHNA auxotroph yet capable of DHNA-mediated EET. We envision that by controlling the expression of a DHNA biosynthesis enzyme (e.g., MenD) on plasmids^29^, the co-culture would allow programmable EET activity for sensing (**Fig. 1**).

We first screened for bacterial candidates qualified to serve as the sender and the receiver in e^-^COSENS. In the sender bacterium, DHNA is biosynthesized from the substrate chorismate through a highly conserved seven-enzyme pathway (MenF, D, H, C, E, B, I) and is subsequently prenylated (MenA + prenyl diphosphate) and demethylated (MenG) to form vitamin K2^30^ (**Fig. 1**). To survey the ubiquitousness of the DHNA biosynthesis pathway, we obtained the identifiers for each of the seven enzymes from the annotated reaction or protein family databases and searched for matched protein entries in the UniProtKB protein sequence database. We identified 2238 phylogenetically distinct bacteria with an intact DHNA biosynthesis pathway (**Fig. 2a, Supplementary Table 1**), indicating that the potential sender candidates are widespread. To survey the potential receiver candidates, we focused on a subset of Firmicutes that were previously predicted to possess a quinone-mediated EET locus^24^. Among the 383 putative EET-capable species in Firmicutes, 324 of them (85%) were found to lack a partial or the entire DHNA biosynthesis pathway (**Fig. 2b, Supplementary Table 2**). This implies that a substantial portion of Firmicutes may depend on exogenous quinones for EET, which qualifies them as potential receiver candidates.

**Fig. 2.**
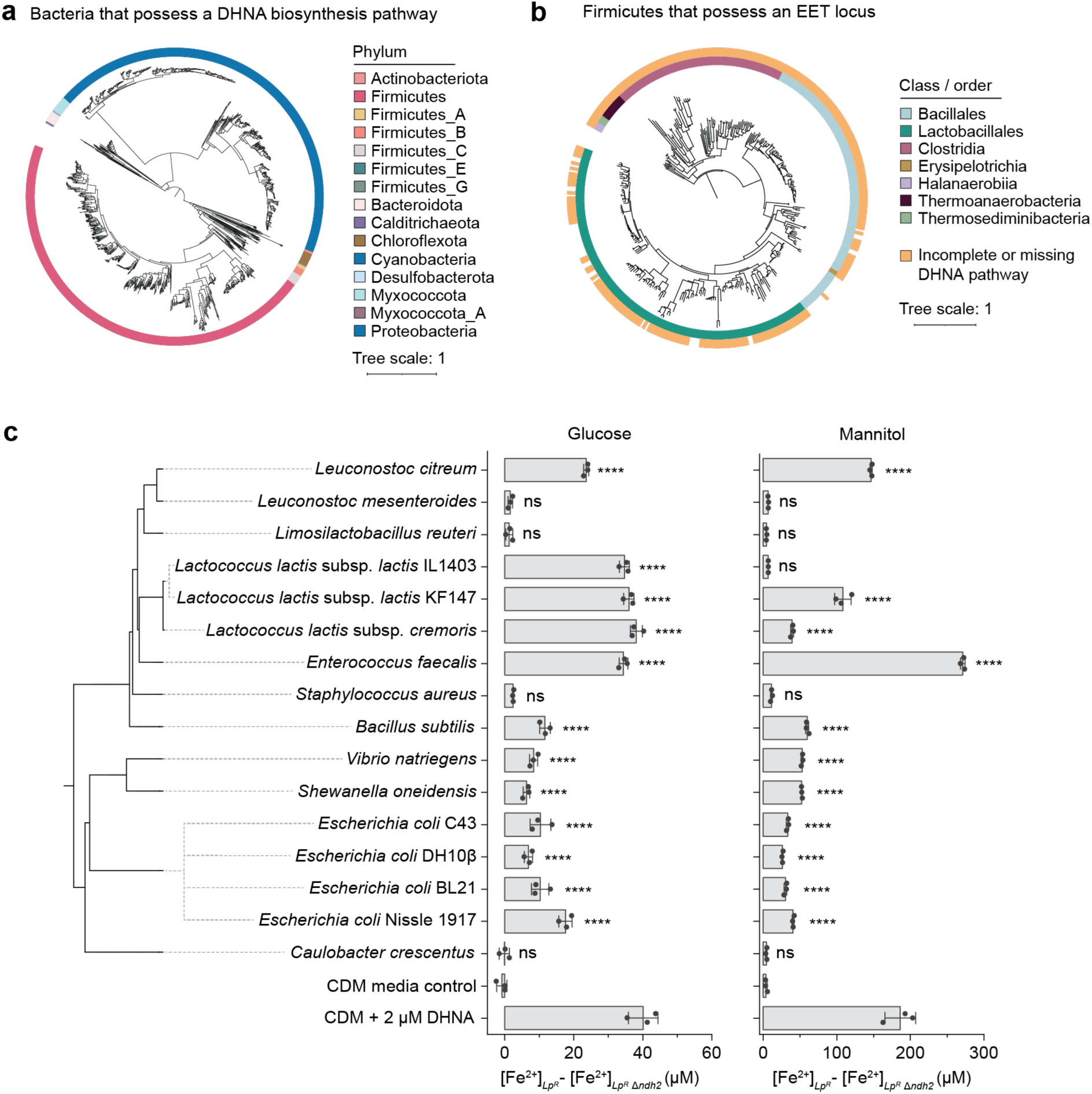
Potential sender and receiver candidates are widespread, and cell-free supernatants from phylogenetically distinct bacteria can activate EET in *L. plantarum.* **a**, phylogenetic tree of bacteria with a complete DHNA biosynthesis pathway. A full list is provided in **Supplementary Table 1**. **b,** Phylogenetic tree of Firmicutes possessing an EET locus^24^ with orange-colored range indicating those with an incomplete or missing DHNA pathway. A full list is provided in **Supplementary Table 2**. **c,** Ndh2-dependent EET in *L. plantarum^R^* can be activated by cell-free supernatants (CFS) from phylogenetically distinct bacteria with either glucose or mannitol as the carbon source. The chemically defined medium (CDM) and CFS from *Caulobacter crescentus* (which lacks the DHNA pathway) serve as negative controls. CDM containing 2 µM DHNA serves as the positive control. Results were reported as the Fe^2+^ concentration reduced by *L. plantarum^R^ ndh2* knockout subtracted from that of *ndh2* wild-type. Data represent mean ± s.d. of three biological replicates. P-values were determined by one-way ANOVA with Tukey’s post hoc test. ****, P < 0.0001 vs. CDM media control; ns, not significant.

We next experimentally validated the sender and receiver candidates. We chose *L. plantarum* as the receiver due to its well-characterized DHNA-mediated EET activity^25,27,28^ and its generally-recognized-as-safe status^26^. Specifically, we used a mutant strain with a *dmkA* knockout (encoding DHNA octaprenyltransferase) to eliminate DHNA consumption and an *ndh1* knockout (encoding NADH:quinone oxidoreductase not essential for EET) to improve electron transfer flux through Ndh2^8^. Hereafter, *L. plantarum* Δ*dmkA*Δ*ndh1* is referred to as “*L. plantarum^R^”* or “*Lp^R^*”. To identify potential senders, we inoculated *L. plantarum^R^* into cell-free supernatant (CFS) collected from 15 putative DHNA-producing bacterial strains across 11 phylogenetically distinct species, as well as a DHNA-null control bacterium (*Caulobacter crescentus*). We also tested glucose and mannitol as carbon sources, as variations in carbon sources can influence quinone synthesis levels^31^. We then quantified the Ndh2-dependent EET by subtracting the iron(III) oxide reduction level of the *L. plantarum^R^*Δ*ndh2* from that of the *ndh2* wild-type. As expected, no EET activity was observed with the CFS from the negative control *C. crescentus* EET. In comparison, EET was activated by the CFS from 12 out of 15 putative DHNA-producing strains when using glucose and by 11 out of 15 strains when using mannitol as the carbon source (**Fig. 2c**). While mannitol supported less bacterial growth compared to glucose (**Extended Data Fig. 1**), the CFS derived from mannitol-based media activated stronger Ndh2-dependent EET (**Fig. 2c**). These results confirm that diverse DHNA-producing bacterial strains can activate EET in the receiver *L. plantarum^R^*, and different carbon sources can modulate the level of EET activity.

### Synthetic co-cultures exhibit electroactivity in a DHNA-mediated Ndh2-dependent manner

Among the bacterial strains whose CFS activated EET in *L. plantarum^R^*, we selected *Lactococcus lactis* subsp*. lactis* KF147 and *E. coli* BL21 as senders to construct co-cultures with *L. plantarum^R^* (**Fig. 3a, d**). *L. lactis* is a Gram-positive, generally-recognized-as-safe bacterium with promise in food and medical applications^32^, and *E. coli* is the Gram-negative model bacterium for synthetic biology. To demonstrate that existing biosensors can be modularly integrated into e^-^COSENS for bioelectronic sensing, we leveraged an *E. coli* BL21 “Marionette” strain that carries a highly optimized genome-integrated sensor array that can respond to diverse small molecules^33^. To enhance DHNA biosynthesis, we eliminated DHNA-consumption pathways by knocking out *menA* (encoding DHNA octaprenyltransferase) in both *L. lactis* and *E. coli*. Additionally, we knocked out *noxAB* (encoding type-II NADH:quinone oxidoreductases) in *L. lactis* to ensure DHNA only interacts with Ndh2 in *L. plantarum^R^*. Hereafter, “*L. lactis^S^*” or “*Ll^S^*” and “*E. coli^S^*” or “*Ec^S^*” imply that these strains contain the respective genomic mutations (**Supplementary Table 3**).

**Fig. 3.**
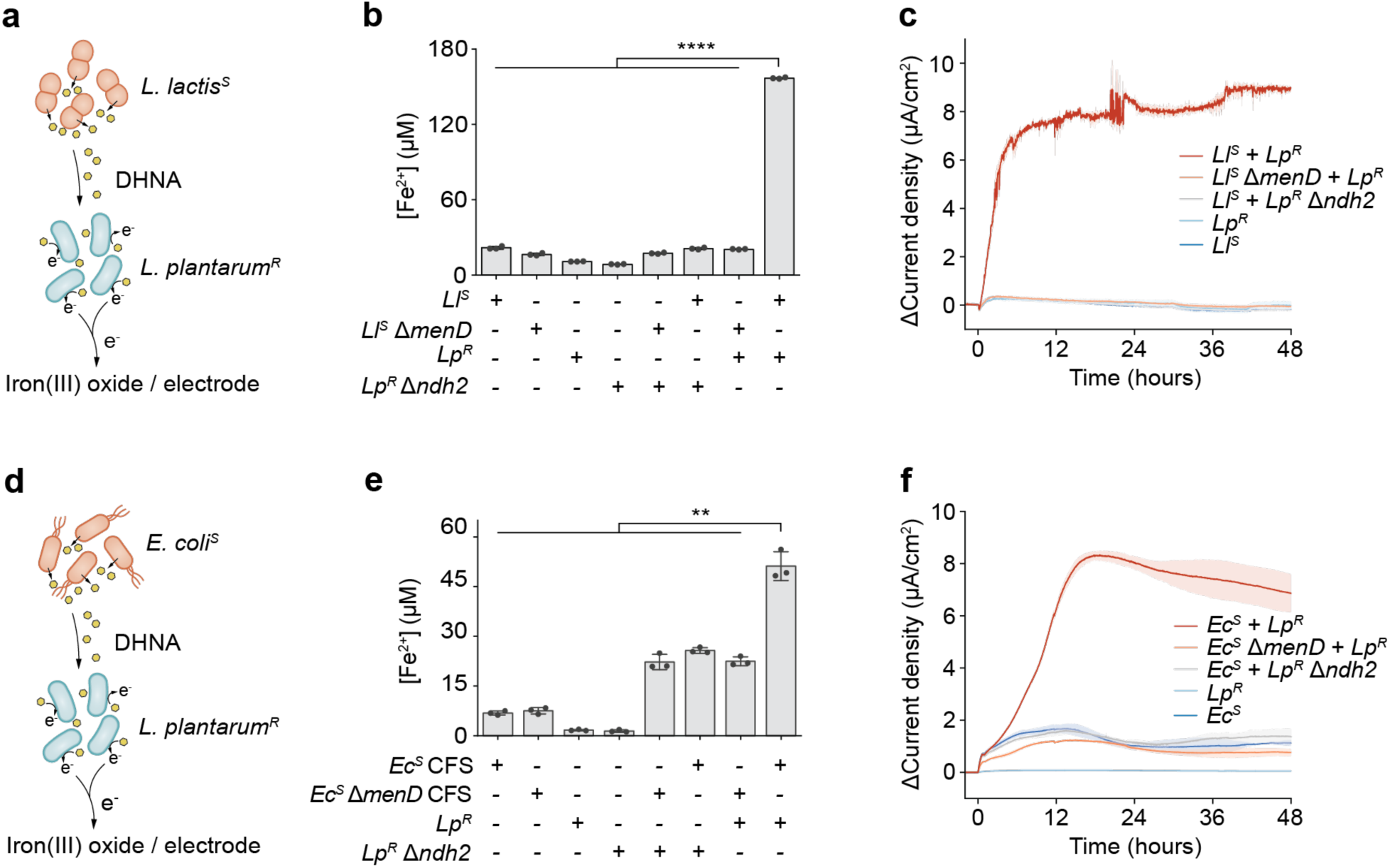
Co-culturing *L. plantarum* with *Lactococcus lactis* or *Escherichia coli* produces electrical signals. a, Schematic showing *Ll^S^-Lp^R^*co-culture reduces extracellular electron acceptors, such as iron(III) oxide or an electrode. **b,c,** Co-cultured *L. lactis^S^* and *L. plantarum^R^* can reduce Fe^3+^ to Fe^2+^ (**b**) and generate current in bioelectrochemical reactors (**c**). Monocultures or co-cultures with mutants deficient in DHNA biosynthesis (*L. lactis* Δ*menD*) and EET (*L. plantarum^R^* Δ*ndh2*) show diminished electroactivity. **d,** Schematic showing *Ec^S^-Lp^R^* co-culture reduces extracellular electron acceptors, such as iron(III) oxide or an electrode. **e,f,** Cell-free supernatant (CFS) from *E. coli^S^* enables *L. plantarum^R^* to reduce Fe^3+^ to Fe^2+^ (**e**), and the co-cultured *E. coli^S^* and *L. plantarum^R^* can produce current in bioelectrochemical reactors (**f**). Monocultures or the mutated *E. coli^S^*Δ*menD* and *L. plantarum^R^* Δ*ndh2* diminish iron reduction and current generation. For experiments in **c** and **f**, co-cultures were injected at 0 h, and the differential (Δ) current density before and after cell injection was reported. Data represent mean ± s.d. of three (**b,e**) or two (**c,f**) biological replicates. P-values were determined by one-way ANOVA with Tukey’s post hoc test. **, P < 0.01; ****, P < 0.0001.

To establish functional co-cultures, we optimized media composition and inoculation ratios. The ideal co-culture conditions should sustain the viability of both sender and receiver strains, support optimal EET activity, and minimize the electrochemical background. We developed mannitol-based minimal media for *Ec^S^-Lp^R^* and *Ll^S^-Lp^R^*co-cultures, respectively (**Supplementary** Fig. 1); these media support the growth of both strains, with mannitol as the carbon source to enhance quinone synthesis and EET. We next determined that a sender-to-receiver ratio of 1:2 to 1:10 enabled maximum fold change in EET using the iron reduction assay (**Extended Data Fig. 2a**). However, *E. coli* was found incompatible with the iron reduction assay due to a high Fe^2+^ background, likely resulting from intracellular iron acquisition during *E. coli* growth^34^. As a result, we initially optimized inoculation ratios using *Ll^S^-Lp^R^* and found these ratios applicable to *Ec^S^-Lp^R^*.

We then tested if co-cultures can reduce iron(III) oxide and produce electrical current via a DHNA-mediated Ndh2-dependent mechanism. When inoculated with iron(III) oxide, monocultures of *L. lactis* and *L. plantarum^R^* and co-cultures deficient in DHNA biosynthesis (*L. lactis^S^* Δ*menD*) or EET (*L. plantarum^R^* Δ*ndh2*) showed no iron reduction activity. Only the functional *Ll^S^-Lp^R^*co-culture reduced a significant amount of iron(III) oxide (**Fig. 3b**). Similarly, when *L. plantarum^R^* was inoculated into CFS from *E. coli^S^*, the iron reduction level lowered if *E. coli^S^*was incapable of DHNA biosynthesis (Δ*menD*) or *L. plantarum^R^* was deficient in EET (Δ*ndh2*) (**Fig. 3e**). These results indicate that both DHNA biosynthesis in *L. lactis* or *E. coli* and EET in *L. plantarum^R^* are essential for extracellular reduction. To test if the co-cultures could produce current, we inoculated both *Ll^S^-Lp^R^*and *Ec^S^-Lp^R^* co-cultures into bioelectrochemical reactors and performed chronoamperometry to monitor current^35^. The co-cultures could transfer electrons to the electrode, resulting in current generation under anaerobic (**Fig. 3c, f**) or microaerobic (**Extended Data Fig. 2b**, c) conditions when both DHNA biosynthesis and EET were functional (**Fig. 3c, f**). While we observed competition between the sender and receiver strains over a 72-hour co-culture period (**Extended Data** Fig. 3), the interspecies interaction did not impair electroactivity. Taken together, these results confirm that sender-receiver co-cultures can exhibit electroactivity, which mechanistically depends on DHNA biosynthesis in *L. lactis^S^* or *E. coli^S^*and Ndh2-dependent EET in *L. plantarum^R^*.

### DHNA-mediated interspecies signaling can be rewired for bioelectronic sensing

Having established electroactive co-cultures, we sought to rewire DHNA-mediated EET to be analyte-dependent for bioelectronic sensing. We reprogrammed the senders to synthesize DHNA only in response to specific environmental stimuli. To achieve this, we knocked out the *menD* gene, which encodes the key regulatory node in the DHNA biosynthesis pathway^29^ from the sender bacterium, and controlled *menD* expression on plasmids under analyte-inducible transcriptional factors (**Fig. 1**). We first tested the well-characterized nisin- and cumate-inducible systems in *L. lactis^S^* and *E. coli^S^*, respectively. Nisin is an antimicrobial peptide produced by *L. lactis* for food preservation that can be sensed by the genomically encoded NisRK two-component system , leading to *menD* transcription from the P_nisA_ promoter (**Fig. 4a**). Cumate is a common inducer molecule that can inhibit CymR^AM^ from repressing the P_cymRC_ promoter^33^, therefore initiating *menD* transcription (**Fig. 4d**).

**Fig. 4.**
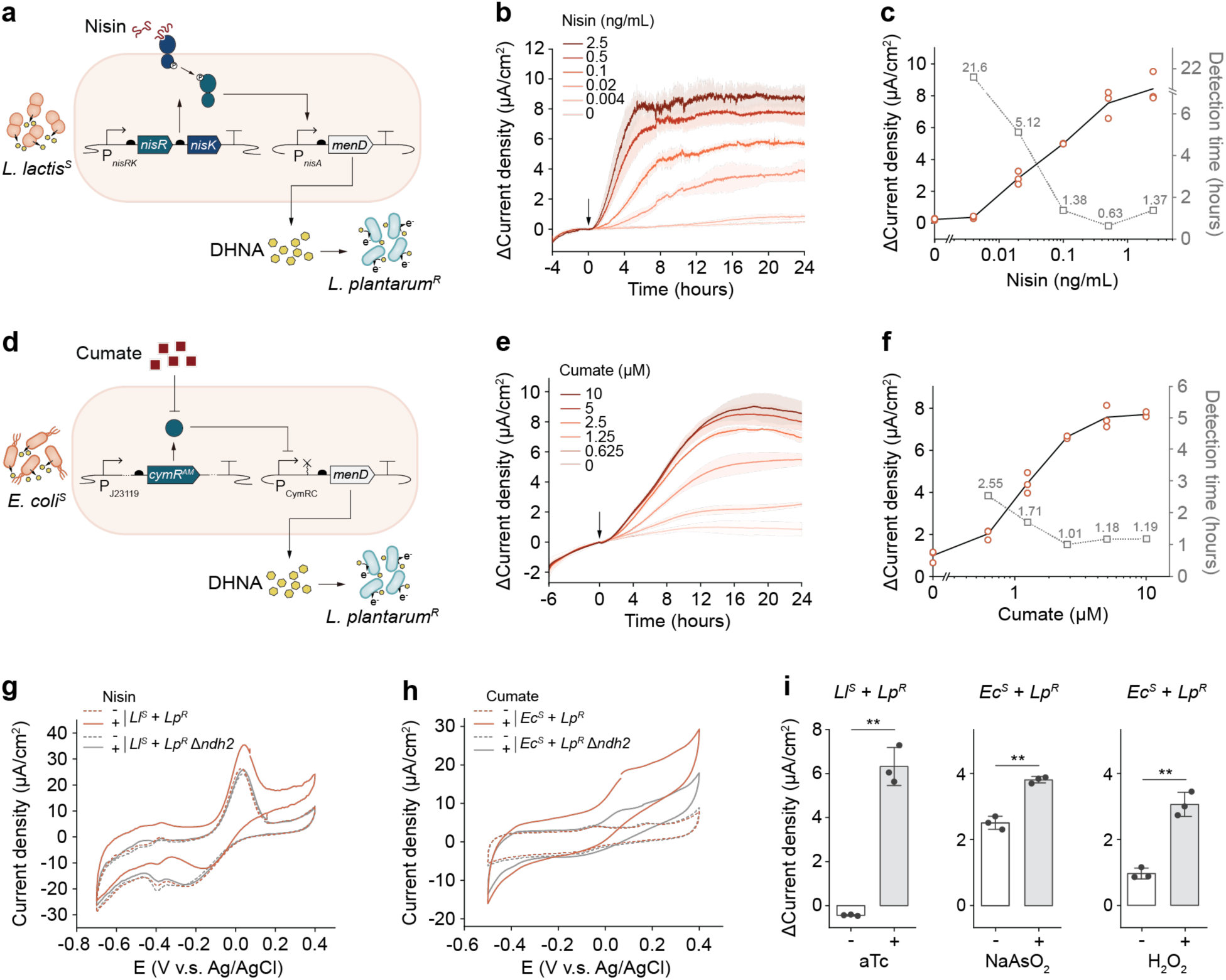
e^-^COSENS allows modular, sensitive, and dose-dependent bioelectronic sensing. **a**, *L. lactis^S^* is engineered to sense nisin and express *menD* for DHNA biosynthesis, which subsequently activates EET in *L. plantarum^R^*to produce electrical signals. **b,** The *Ll^S^*-*Lp^R^*co-culture produces current upon nisin induction at 0 h (indicated by the arrow). The differential (Δ) current densities before and after nisin induction are proportional to the nisin concentrations (0.004-2.5 ng/mL). **c,** Quantitative analysis of Δcurrent density and the 95% confidence time of detection (P < 0.05 compared to no induction control) as a function of nisin concentration. **d,** *E. coli^S^* is engineered to sense cumate and express *menD* for DHNA biosynthesis, which subsequently activates EET in *L. plantarum^R^*to produce electrical signals. **e,** The *Ec^S^*-*Lp^R^*co-culture produces current upon cumate induction at 0 h (indicated by the arrow). The differential (Δ) current densities before and after nisin induction are proportional to the cumate concentrations (0.1-10 µM). **f,** Quantitative analysis of Δcurrent density and 95% confidence time of detection (P < 0.05 compared to no induction control) as a function of cumate concentration. **g,h,** Cyclic voltammetry (CV) analysis revealing signal amplification capacity of e^-^COSENSE (red curves) compared to non-EET conditions (Δ*ndh2*, gray curves). CV was conducted at peak current post-nisin or cumate induction. Dotted curves represent no induction controls. **i,** Reprogram e^-^ COSENS to sense anhydrotetracycline (aTc), arsenite (NaAsO2), or hydrogen peroxide (H2O2). The results show Δcurrent density at 12 h post-induction. Time-dependent data and plasmid designs can be found in **Extended Data 6**. Data in **g** and **h** are representative results of three biological replicates. All other data represent mean ± s.d. of three biological replicates. P-values were determined by a two-tailed unpaired t-test. **, P < 0.01.

We optimized the genetic circuits for maximal sensing fold change and evaluated e^-^COSENS’s sensing capability in electrochemical reactors. To reduce basal expression, we screened degenerate ribosome binding sites (RBSs) with varying translation initiation rates (TIRs) for *menD* expression (**Extended Data Fig. 4a**, d). RBSs with the lowest TIRs yielded the largest fold changes in iron reduction in response to nisin (**Extended Data Fig. 4b**, c) or cumate (**Extended Data Fig. 4e**, f) and were thus selected for subsequent use. We then inoculated co-cultures in electrochemical reactors and monitored current generation upon nisin or cumate induction. As controls, co-cultures with the empty-vector senders or EET-deficient *L. plantarum^R^* had no response to inducers (**Extended Data Fig. 4g**, h). The functional *Ll^S^-Lp^R^* (**Fig. 4b**) and *Ec^S^-Lp^R^* (**Fig. 4e**) co-cultures responded to inducers and generated differential (Δ) current densities proportional to the inducer concentrations (**Fig. 4c, f**). The time required to significantly differentiate (P<0.05) sensing signals from the background was negatively proportional to the inducer concentrations, with the shortest time being 0.63 and 1.01 hours for nisin and cumate sensing, respectively (**Fig. 4c, f**). These results indicate that by controlling DHNA-mediated EET, synthetic co-cultures can be rewired for quantitative bioelectronic sensing, and the sensing ability necessitates both DHNA biosynthesis and EET functionalities.

Bioelectronic sensors have also used voltammetric techniques to detect redox-active signaling molecules like DHNA^37–40^; this led us to compare the signal intensity from direct voltammetric measurement of DHNA with that from DHNA-mediated EET. To do so, we compared cyclic voltammetry (CV) profiles of co-cultures comprising EET-capable or deficient (Δ*ndh2*) *L. plantarum^R^*after nisin or cumate induction. When EET was deficient, CV was able to detect a DHNA catalytic wave around a midpoint of -25 mV (vs. Ag/AgCl) for *E. coli^S^* but not for *L. lactis^S^* (**Fig. 4g, h**). However, when EET was functional, such a catalytic wave became detectable for *L. lactis^S^* and was significantly more prominent for *E. coli^S^* (**Fig. 4g, h**). At 200 mV (the potential for chronoamperometry), the current density was 2.0-or 3.3-fold higher for EET-capable *Ec^S^-Lp^R^* or *Ll^S^-Lp^R^* co-cultures, respectively, compared to deficient controls. This increase in current could result from continuous redox cycling of DHNA between the electrode and *L. plantarum*’s EET pathway, which amplifies the electrical signals. Thus, e^-^COSENS offers advantages over direct CV detection by providing signal amplification and greater sensitivity.

As many environmental molecules are redox-active, we asked whether these background molecules would interfere with DHNA-mediated sensing. We sequentially exposed co-cultures to 250 nM microbe- or plant-derived redox-active molecules, including heme, pyocyanin, phenazine-1-carboxylic acid, riboflavin, flavin adenine dinucleotide, flavin mononucleotide, and phylloquinone. Although some background current was observed and the electroactivity of *Ll^S^-Lp^R^* was slightly impaired, these redox-active molecules did not affect current generation in response to nisin or cumate induction (**Extended Data Fig. 5a**, b). Additionally, we tested the influence of environmental DHNA and observed that 50 nM DHNA induced a high background current, which inhibited nisin sensing in *Ll^S^-Lp^R^* (**Extended Data Fig. 5c**). However, cumate sensing in *Ec^S^-Lp^R^* was not affected despite the DHNA background current (**Extended Data Fig. 5d**). The repression might be due to exogenous DHNA allosterically inhibiting MenD with a half-maximal inhibitory concentration varying from about 50 nM to 4 µM in different bacterial species^41,42^, which would suppress endogenous DHNA biosynthesis (**Extended Data Fig. 5e**). These results suggest that, while exogenous DHNA influences e^-^COSENS to varying extents depending on the sender bacterium, e^-^COSENS is generally resilient against many common redox-active molecules.

Having shown that e^-^COSENS can detect common inducers, we sought to probe its modularity to detect other analytes, including anhydrotetracycline (aTc, a derivative of the antibiotic tetracycline), arsenite (NaAsO_2_, a heavy metal contamination in water), and hydrogen peroxide (H_2_O_2_, an inflammation biomarker). We used established transcriptional systems: *tetR*-P*_xyl/2xtetO_* for aTc sensing in *L. lactis*^43^, *arsR*-P*_arsOC2_* for arsenite sensing in *E. coli*^44^, and *oxyR*-P*_oxyS_* for H_2_O_2_ sensing in *E. coli*^45^ (**Extended Data Fig. 6a**, c, e). With optimized RBSs, we detected significant iron reduction at 1.3-10 µM aTc, 0.01-10 µM arsenite (the World Health Organization limit of 10 ppm or ∼0.1 µM in drinking water), and 1-100 µM H_2_O_2_ (**Extended Data Fig. 6a**, c, e). When tested in bioelectrochemical reactors, both *Ll^S^-Lp^R^*and *Ec^S^-Lp^R^* produced current in response to the respective analytes (**Fig. 4i, Extended Data** Fig. 6b, d, f). However, a relatively high background was observed for arsenite and H_2_O_2_ sensing circuits, suggesting further optimization is needed to reduce basal expression. Nonetheless, these results highlight e^-^COSENS’s unprecedented modularity for bioelectronic sensing, allowing flexible switching of the sender bacterium and associated genetic circuits to detect diverse analytes.

### e^-^COSENS functions in diverse environments and within microbial communities

We next explored whether e^-^COSENS could be applied in real-world environments. As *L. lactis* and *L. plantarum* are GRAS species commonly used for food and health applications^32,46^, we tested if their co-culture could sense aTc in milk and produce electrical signals, showing potential for food quality monitoring. Engineered *E. coli* modalities have shown promise for applications in various environments and mammalian hosts^5,47–49^; we examined whether its co-culture with *L. plantarum* could be applied for pollutant and biomarker monitoring by testing arsenite and H_2_O_2_ bioelectronic sensing in bayou water and artificial saliva. Milk was obtained from a local grocery store, and bayou water was collected from the Brays Bayou in the Houston area. Artificial saliva mimics natural saliva’s chemical and physical characteristics^50^ and was used as a substitute for saliva to enable testing in larger quantities. These samples have distinct physicochemical properties, varying in total organic carbon (TOC), pH, electrical resistance, and light transmittance (**Fig. 5a**). Notably, milk exhibited 0% transmittance for light at 509 nm (1 cm path length), which is the emission wavelength of green fluorescent protein (GFP), making it an opaque environment where GFP-based biosensors would be ineffective. Moreover, the samples represent a carbon source level varying over a factor of 10^4^, with TOC of 5×10^4^ mg/L for milk to 5 mg/mL for bayou water.

**Fig. 5.**
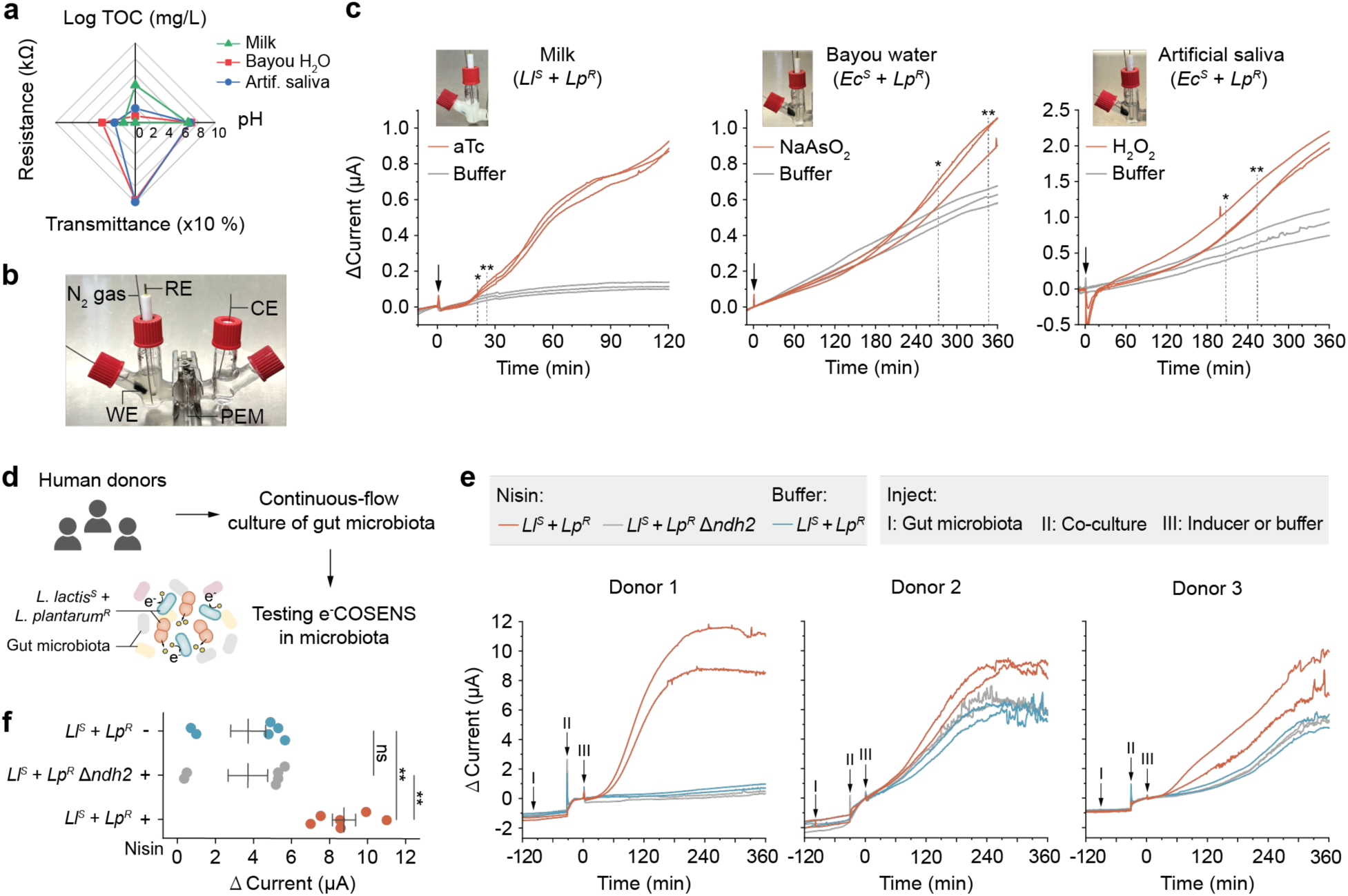
e^-^COSENS enables sensing in food, environmental, and human-relevant environments. **a**, Physicochemical properties of milk, bayou water, and artificial saliva samples. TOC, total organic carbon. Transmittance was determined at 509 nm, the emission wavelength of green fluorescent protein. **b,** Photo of the 5 mL-volume bioelectrochemical reactors. N2 gas, nitrogen gas; WE, working electrode; CE, counter electrode; RE, reference electrode; PEM, proton exchange membrane. **c,** The *Ll^S^*-*Lp^R^* co-culture detects aTc (5 ng/mL) in milk, and the *Ec^S^*- *Lp^R^* co-culture detects NaAsO2 (2.5 μM) and H2O2 (25 μM) in bayou water and artificial saliva, respectively. The arrows indicate analyte injection. The differential (Δ) current before and after analyte addition was reported. The asterisks indicate 95% (*, P < 0.05) or 99% (**, P < 0.01) confidence time of detection. Data represent three biological replicates. **d,** Testing *Ll^S^*-*Lp^R^*co-culture in the gut microbiota from three human donors. The large intestinal microbiota was cultivated in continuous-flow minibioreactor arrays^53^. **e,** *Ll^S^*-*Lp^R^*co-culture can sense nisin (2.5 ng/mL) and produce distinguishable current within the three gut microbiota samples. Arrows I, II, and III indicate gut microbiota, co-culture, and nisin/buffer injection. **f,** Statistical analysis of the current levels in **e** at 360 min. Data represent mean ± s.e.m. of the six biological replicates obtained from three donors. P-values were determined by two-tailed unpaired t-test (**c**) or one-way ANOVA with Tukey’s post hoc test (**f**). *, P < 0.05; **, P < 0.01; ns, not significant.

To evaluate e^-^COSENS’s functionality in these environments, we inoculated the exponential-phase *Ll^S^-Lp^R^* or *Ec^S^-Lp^R^* co-cultures into bioelectrochemical reactors containing 5 mL samples spiked with respective analytes (**Fig. 5b**). The *Ll^S^-Lp^R^* co-culture detected aTc (5 ng/mL) in milk with 95% and 99% confidence within 21 and 25.8 minutes (**Fig. 5c**). Despite a higher background, the arsenite (2.5 µM) was detected by *Ec^S^-Lp^R^*co-culture with 95% and 99% confidence in 273 and 347 minutes (**Fig. 5c**). Similarly, the H_2_O_2_ (25 µM) could also be sensed by *Ec^S^-Lp^R^* co-culture with 95% and 99% confidence in 207 and 253 minutes (**Fig. 5c**). The detection time varied across different environments and co-cultures, potentially affected by the basal current level, the nutrient availability, and the cell viability (**Extended Data Fig. 7a**). The fastest response was achieved in the nutrient-rich milk sample (**Fig. 5c**) with the low basal expression aTc-sensing genetic circuit (**Extended Data Fig. 6a**, b). Moreover, no differential current was observed if *L. lactis^S^* or *E. coli^S^* carried empty vectors or if *ndh2* was knocked out from *L. plantarum^R^*(**Extended Data Fig. 7b**-g), indicating that sensing relied on functional co-cultures. These results demonstrate that 1) electrical signals can be detected in opaque environments (milk) where optical biosensors are inoperable; 2) the universal receiver *L. plantarum^R^* is applicable in distinct environments, which is not surprising given *L. plantarum*’s nomadic lifestyle and adaptability to diverse habitats^51^; 3) by pairing different senders with *L. plantarum^R^*, e^-^COSENS can be tailored for bioelectronic sensing in diverse environments. Since *L. lactis* and *L. plantarum* are present in the human gastrointestinal tract^52^, we additionally asked whether their co-culture can be used for bioelectronic sensing within the human gut microbiota. Using continuous-flow minibioreactor arrays (MBRAs)^53^, we cultivated microbiota from the large intestine contents of three distinct human donors. We then inoculated each microbiota along with *Ll^S^-Lp^R^* co-culture into bioelectrochemical reactors to test for nisin sensing (**Fig. 5d**). The co-culture survived (**Extended Data Fig. 8a**-c) and detected nisin in all three microbiota samples, producing current distinguishable from the EET-deficient and buffer controls (**Fig. 5e, f**). Donor 1 exhibited the lowest background current, while donors 2 and 3 showed a moderately high background (**Fig. 5e**). The variations in background levels can be attributed to the different microbiota composition across the three donors (**Extended Data Fig. 8d**). These results suggest that e^-^COSENS is effective within complex microbial communities despite potential interference from neighboring microbes.

### A miniature bioelectronic device enables portable signal detection

Prior electrical signal detection was achieved using glassy electrochemical reactors and high-precision potentiostat (**Methods**); these instruments are cumbersome, high-cost, and have very limited portability. To enable low-cost, portable, and energy-friendly bioelectronic sensing, we developed a miniature clay-based microbial fuel cell (MFC) device for detecting electrical signals from e^-^COSENS (**Fig. 6a**, **Extended Data Fig. 9a**). The device is fabricated by sandwiching a proton exchange membrane (PEM) between a screen-printed Ag/AgCl cathode and a carbon anode stacked with a 6 mm carbon felt round (**Fig. 6a**). Particularly, the PEM was exfoliated from natural vermiculite clay, showing high biocompatibility and extraordinary ion conductivity of up to 2×10^-2^ S/cm^54^. Manufactured in Boston and shipped to Houston for testing, the devices maintained good performance, demonstrating chemical and mechanical stability. Moreover, a single device costs less than three dollars and can be manufactured in a high-throughput way (**Extended Data Fig. 9b**).

**Fig. 6.**
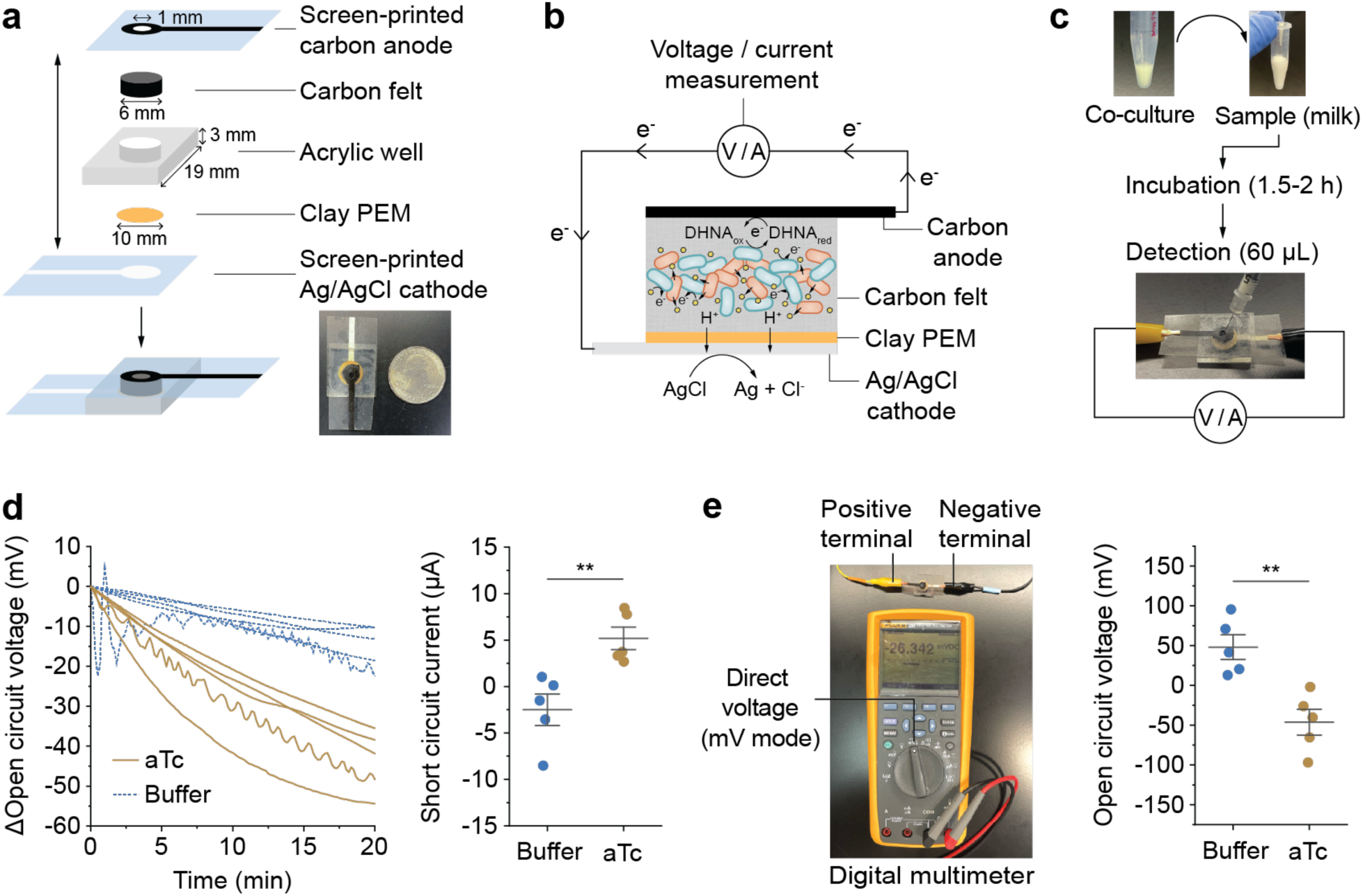
A miniature bioelectronic device simplifies signal detection. **a**, The assembly schematic of the clay-based microbial fuel cell and a comparison of the size between the device and a coin. PEM, proton exchange membrane. **b,** Electrons generated through the DHNA-mediated EET in e^-^COSENS can accumulate in carbon felt or move from the carbon anode to the Ag/AgCl cathode via an external circuit, allowing for open circuit voltage (OCV) or short circuit current (SCA) measurements. At the cathode, AgCl is reduced, and H^+^ movement through the clay PEM maintains charge neutrality. **c,** The workflow of signal detection. Co-culture was inoculated into the sample or media and incubated for 1.5-2 hours, and a 60 µL portion was then injected into the device for OCV or SCA measurements. **d**, OCV and SCA measurements for aTc detection in milk. Differential (Δ) OCV was reported by subtracting OCV at t = 0 from subsequent OCV values. SCA was determined by linear sweep voltammetry at voltage = 0 vs. OCV. **e,** OCV measurement using a digital multimeter. The positive terminal of the multimeter is connected to the carbon anode of the device, and the negative terminal is connected to the Ag/AgCl cathode. OCV measured by the digital multimeter enables aTc detection in milk. Data represent five biological replicates across two independent experiments on different days. Error bars in **d, e** represent mean ± s.e.m. P-values were determined by a two-tailed unpaired t-test. **, P < 0.01.

We detected electrical signals from e^-^COSENS using the clay-based MFC device by measuring the open circuit voltage (OCV) and short circuit current (SCA) (**Fig. 6b**). The OCV measures the potential difference between anode and cathode; a faster drop of OCV is anticipated for DHNA-mediated EET because DHNA reduction accumulates electrons on the anode, leading to a more rapid decrease in anodic potential compared to the EET-deficient control. When the circuit is closed, EET would enable more electron transfer from the anode to the cathode, resulting in higher SCA. To prepare for detection, the exponential-phase co-cultures were incubated for 1.5-2 hours to allow DHNA secretion. Then, 60 µL of the co-cultures were injected into the acrylic well of the device for OCV and SCA measurements (**Fig. 6c**). As expected, we observed a more rapid drop of OCV and a higher SCA for EET-capable *Ll^S^-Lp^R^*and *Ec^S^-Lp^R^* co-cultures when compared to those containing *L. plantarum^R^* Δ*ndh2* (**Extended Data Fig. 9c**, d). Consistently, when tested with aTc sensing in milk with *Ll^S^-Lp^R^*(**Fig. 6c**), the faster drop of OCV and higher SCA reflected the detection of aTc (**Fig. 6d**). These results demonstrate that the mini bioelectronic device supports signal detection from e^-^COSENS using microliter-scale samples.

We also demonstrated using a digital multimeter to measure electrical signals (**Fig. 6e**). By connecting the positive terminal of the multimeter to the carbon anode of the device and the negative terminal to the Ag/AgCl cathode, the multimeter displayed lower OCV values for EET-capable co-cultures compared to deficient controls (**Extended Data Fig. 9e**) and also reflected aTc sensing (**Fig. 6e**). This highlights the potential of e^-^ COSENS for portable bioelectronic sensing with a digital multimeter as a simple and accessible method for electrical signal readout.

## Discussion

In this study, we developed e^-^COSENS, a modular bacterial co-culture platform for designing bioelectronic sensors. This system consists of a “sender” bacterium that produces DHNA in response to target analytes and a “receiver” bacterium that generates electrical signals through DHNA-mediated EET. We showcased the versatility of e^-^ COSENS by co-culturing two phylogenetically distinct senders, *L. lactis* and *E. coli,* with the universal receiver *L. plantarum*. By leveraging genetic sensing circuits functional in either *L. lactis* or *E. coli*, we demonstrated the detection of nisin, aTc, cumate, arsenite, and H_2_O_2_ in lab-based media, milk, bayou water, artificial saliva, and the human microbiome. Moreover, we designed a miniature MFC device for low-cost, low-energy, and portable signal detection using a commercial digital multimeter.

The modularity of e^-^COSENS overcomes several key bottlenecks for bioelectronic sensor development, including limited availability of chassis and complex engineering strategies. First, it removes the need to tediously engineer electroactive bacteria to create bioelectronic sensors. Instead, a wide range of non-electroactive bacteria that produce DHNA (**Fig. 2a, c**) can be potentially coupled with a receiver bacterium for standardized electrical signal conversion. Second, it significantly broadens the range of detectable analytes by allowing flexible selection of the sender bacterium and associated genetic sensing elements^55^. This includes microbial chassis with large genetic toolboxes, such as but not limited to *E. coli*^33,56^, *L. lactis*^32,57^, *Vibrio natriegens*^58^*, Bacillus subtilis*^59^ (**Fig. 2c**). Moreover, e^-^COSENS opens up new environments where bioelectronic sensors can be deployed. With the generally-recognized-as-safe *L. lactis* and *L. plantarum*, we demonstrated efficient use of bioelectronic sensors in food products and human-relevant contexts (**Fig. 5**). This approach could possibly extend to other contexts by pairing the multi-habitat-adaptable receiver *L. plantarum*^60^ with appropriate senders, such as *V. natriegens* for sensing in marine environments^61^ or *B. subtilis* for soil and plant applications^62^ (**Fig. 2c**). With these advantages, e^-^COSENS could allow the creation of customized bioelectronic sensors for specific applications.

We envision several improvements to further advance the e^-^COSENS platform. To address environmental DHNA’s attenuation on e^-^COSENS inducibility (**Extended Data Fig. 5c**, d), likely due to its allosteric inhibition on MenD (**Extended Data Fig. 5e**), a DHNA-insensitive MenD mutant can be created to eliminate this inhibitory effect^41,42^. To minimize background electrical signals caused by environmental DHNA (**Extended Data Fig. 5c**, d) and other microbes in the community (**Fig. 5e**), the co-culture can be encapsulated in a hydrogel matrix^5,63^, which would enrich the sensing signals around the electrodes, reducing the impact of noise and enhancing the signal-to-noise ratio. Additionally, carbon source availability and type may also impact sensor performance (**Fig. 2c and Extended Data Fig. 1**). When applying e^-^COSENS in the field, carbon sources can be provided during sample incubation (**Fig. 6b**) or incorporated along with an encapsulation matrix. To stabilize DHNA biosynthesis and sensor output across different carbon sources, the putative carbon source-sensitive promoter of the *men* operon^64^ can be genomically replaced with a constitutive promoter to allow consistent transcription of the *men* operon.

Looking forward, the e^-^COSENS platform offers opportunities to combine biological and electronic systems for new sensing technologies. Of particular interest is e^-^COSENS’s ability to generate electrical energy during sensing, which can enable self-powered bioelectronic sensors ideal for long-term and remote monitoring^65^. With endeavors in electronic device miniaturization (**Fig. 6**), e^-^COSENS can be incorporated into various compact electronic modalities, such as paper-based electrochemical systems^10,11,16^ or ingestible electronic capsules^48,49^. These bacterial-electronic hybrid systems will offer valuable tools for effective monitoring of environment, food, and health parameters.

## Methods

### Strains

A list of strains is provided in **Supplementary Table 3**. *Escherichia coli* DH5α (NEB) was used for all molecular cloning. The receiver strain *Lactiplantibacillus plantarum* NCIMB8826 Δ*dmkA*Δ*ndh1,* and the EET null *L. plantarum* Δ*dmkA*Δ*ndh1*Δ*ndh2* have been previously reported^8^.

To develop the sender *L. lactis^S^*, the parental strain *Lactococcus lactis* subsp*. lactis* KF147 was genome-modified to knock out *menA*, *noxAB*, and *menD* using double-crossover homologous recombination. Upstream and downstream homologous arms (about 600 base pairs each) flanking the targeted gene were inserted into the NotI-EcoRI digested suicide plasmid pRV300^66^. The resulting plasmids were electroporated into *L. lactis*, and genome-integrated mutants were selected on 0.5% glucose M17 (gM17) agar plates containing 5 µg/mL erythromycin at 30 °C and subsequently verified by colony PCR. A single genome-integrated colony was inoculated in 3 mL gM17 without antibiotics at 30 °C and passaged daily at a 1:1000 dilution. Starting from day 6, cells were spread on gM17 plates without antibiotics, and the following day, replica-plated on gM17 agar plates containing 5 µg/mL erythromycin. Erythromycin-sensitive colonies were picked, and gene deletion was confirmed by colony PCR and DNA sequencing.

To develop the sender *E. coli^S^*, the *Escherichia coli* BL21 “Marionette” strain with a “sensor array” integrated into the genome^33^ was used as the parental strain and was genome-modified to knock out *menA* and *menD* using CRISPR-Cas9-assisted genome editing^67^. The donor DNA containing upstream and downstream homologous arms (about 500 base pairs each) flanking the targeted gene was synthesized as a linear double-stranded DNA fragment (Twist Bioscience). Single-guide RNA (sgRNA) specific to the targeted gene was introduced into the pTargetF backbone (Addgene, #62226). The *E. coli* strain harboring pEcCas (Addgene, #73227) was grown overnight and then subcultured in 25 mL lysogeny broth (LB) containing 50 µg/mL kanamycin with an initial OD_600_ = 0.05. When OD_600_ reached 0.1, 40 mM arabinose was added to induce λ-Red recombinase expression. Cells were harvested at an OD_600_ of 0.4-0.6 by centrifugation and made electrocompetent. Competent cells (50 µL) were electroporated with 400 ng linear donor DNA and 100 ng pTargetF (containing gene-specific sgRNA). Colonies were selected on LB agar plates containing 50 µg/mL kanamycin and 50 µg/mL spectinomycin at 37 °C. Positive mutants were then verified by colony PCR and DNA sequencing. pTargetF and pEcCas were cured by growing cells in the presence of 10 mM rhamnose or 5 g/L glucose and 10 g/L sucrose, respectively, according to the previous study^67^.

### Culture conditions

All molecular cloning experiments were performed in LB or Terrific Broth. To prepare *Ll^S^*- *Lp^R^* co-cultures, *L. plantarum^R^* was initially grown in 3 mL commercial MRS (deMan, Rogosa, and Sharpe) broth at 37 °C without shaking for 15 h, and an aliquot was subcultured (1:100 v/v) into a desired volume of 1% mannitol MRS broth (mMRS) (**Supplementary Table 8**) at 37 °C without shaking for 15 h. *L. lactis^S^* was initially grown in 3 mL 0.5% glucose M17 (gM17) at 30 °C without shaking for 15 h, and an aliquot was subcultured (1:100 v/v) in a desired volume of 1% mannitol M17 (mM17) at 30 °C without shaking for 15 h. For iron reduction assay, evaluation of co-culture electroactivity, and bioelectronic sensing in lab-based media, the stationary-phase cultures of *L. lactis^S^* and *L. plantarum^R^* were washed twice in 1X phosphate-buffered saline (PBS) and resuspended in a blended medium containing 1% mannitol chemically defined medium (mCDM) (**Supplementary Table 6**) and mM17 in a 20:1 volume ratio. This medium supported the growth of *L. lactis^S^*for sensing while maintaining a low electrochemical background. Co-cultures were created by mixing the two resuspended strains at the indicated OD_600_. For bioelectronic sensing in milk and gut microbiota samples, stationary-phase *L. lactis^S^*and *L. plantarum^R^* were subcultured in mM17 or mMRS, respectively, at an initial OD_600_ of 0.1 until reaching the exponential phase (OD_600_ = 0.4-0.6). Cells were then washed once with PBS and mixed in mM17 at the indicated OD_600_ for injection into samples. When *L. lactis^S^* or *L. plantarum^R^* carried erythromycin resistance plasmids, all media and samples above (except gut microbiota samples) were supplemented with 10 µg/mL erythromycin.

To prepare *Ec^S^*-*Lp^R^* co-culture, *L. plantarum^R^* was grown following the same procedure described above. *E. coli* was initially grown in 3 mL LB at 37 °C with 200 rpm shaking for 15 h, and an aliquot was subcultured (1:100 v/v) in a desired volume of M9 minimal medium containing 1% mannitol (minimal mM9) (**Supplementary Table 7**) at 37 °C with 200 rpm shaking for 15 h. To evaluate co-culture electroactivity and bioelectronic sensing in lab-based media, stationary-phase cultures of *E. coli^S^* and *L. plantarum^R^* were washed twice in 1X PBS and resuspended in minimal mM9. Co-culture was created by mixing the two resuspended strains at the indicated OD_600_. For bioelectronic sensing in bayou water and artificial saliva samples, stationary-phase *E. coli* and *L. plantarum^R^* were subcultured in minimal mM9 or mMRS at an initial OD_600_ of 0.05 or 0.1, respectively, until reaching the exponential phase (OD_600_ = 0.4-0.6). Cells were then washed once with PBS and mixed at the indicated OD_600_ in minimal mM9 (2.5% mannitol) for injection into samples. When *E. coli^S^*or *L. plantarum^R^* carried chloramphenicol resistance plasmids, 34 µg/mL or 10 µg/mL chloramphenicol was added to the monoculture medium of *E. coli^S^* or *L. plantarum^R^*, respectively. Due to the redox activity of chloramphenicol, a low concentration of 3 µg/mL was used in the co-culture medium and samples for electrochemical analysis.

### Plasmid construction

A list of plasmids is provided in **Supplementary Table 4**. The sequence of genetic parts is provided in **Supplementary Table 5**. Plasmids for *L. lactis^S^* were constructed using a backbone derived from the shuttle vector pECGMC3 (Addgene #75441). For nisin sensing, the NisKR two-component system is natively expressed in *L. lactis* KF147^60^. The P*_nisA_* promoter and ribosome binding site (RBS) were synthesized as a DNA fragment (Twist Bioscience) based on the sequence from pNZ8048^68^, and cloned upstream of the *menD* gene amplified from the genome of *L. lactis* KF147 using Golden Gate assembly^69^. To tune RBS strength, a 30 bp sequence upstream of the *menD* start codon was designed to contain four degenerate nucleotides in RBS (NNGGNGN), and an RBS library with varying translation initiation rate (TIR) was generated using the RBS library Calculator^70,71^. Four RBS variants with TIR ranging from 10^2^ to 10^4^ were selected for experimental testing. For aTc sensing, the *tetR*-P*_tetR_*_-D7_-P*_xyl_*_/2x*tetO*_ cassette^43^ was synthesized as a DNA fragment (Twist Bioscience). The same RBS library used for P*_nisA_* was placed after P*_xyl_*_/2x*tetO*_ to optimize *menD* expression.

Plasmids for *E. coli^S^* were constructed using a medium-copy backbone containing a chloramphenicol-resistant gene and p15A origin. For cumate sensing, the P_CymRC_ promoter^33^, SarJ insulator^33^, a synthetic RBS designed by RBS Calculator^70,71^, and the *menD* gene amplified from the genome of *E. coli* BL21 were integrated into the backbone using Golden Gate assembly. A degenerate RBS (NNGGNGN) library was designed, and four RBS variants (TIR ranging from 10^1^ to 10^3^) were selected to optimize *menD* expression. For arsenite and hydrogen peroxide sensing, the P_J109_-RBS_30_-*arsR*-DT54- P*_arsOC2_*^44^ and P*_proD_*-RBS*_oxyR_*-*oxyR*-DT54-P*_oxyS_*^45^ cassettes were synthesized and assembled with SarJ and a weak RBS from the degenerate RBS library to drive *menD* expression. DT54^56^ is a double terminator that prevents cross-transcription. To optimize arsenite sensing, three synthetic RBSs (TIR = 10^2^-10^3^) were tested to replace RBS_30_ and tune *arsR* expression.

Plasmids for *L. plantarum^R^* were constructed using a backbone derived from pSIP403^72^. An ampicillin-resistant gene *ampR* was inserted into the backbone for molecular cloning. The erythromycin-resistant gene *ermR* was replaced by the chloramphenicol-resistant gene *cat* for co-culture with *E. coli^S^*.

### Bioinformatic and phylogenetic analysis

Identifiers for the seven enzymes in the DHNA biosynthesis pathway were obtained from the annotated reaction database Rhea^73^, as well as the protein family databases InterPro^74^ and NCBIfam^75^. A list of identifiers for each enzyme is listed in **Supplementary Table 10**. These identifiers were used to search for matching protein entries in the protein sequence database UniProtKB (assessed on September 18, 2023). The NCBIfam identifier (TIGR00369) for *menI* (1,4-dihydroxy-2-naphthoyl-CoA hydrolase) also includes other hotdog fold thioesterases, and protein entries were filtered to retain those specifically annotated as 1,4-dihydroxy-2-naphthoyl-CoA hydrolase/thioesterase. The resulting protein entries were sorted by organism name using an in-house MATLAB program. The taxa of organisms with no missed genes in the DHNA biosynthesis pathway were obtained from the Genome Taxonomy Database^76^ (assessed on November 8, 2023), and the phylogenetic trees were visualized using iTOL (https://itol.embl.de/). To identify potential EET-capable Firmicutes with incomplete DHNA biosynthesis pathways, previously identified Firmicutes harboring the EET locus^24^ were compared with organisms with no missed DHNA biosynthesis genes.

### Iron reduction assays for cell-free supernatants or co-culture

*L. plantarum^R^* and *L. lactis^S^* were resuspended in the mCDM:mM17=20:1 medium to OD_600_ of 2 or 0.2, respectively. In a 96-deep well plate, a 100 µL portion of each cell resuspension was mixed along with 200 µL of reagent mixture containing 4 mM iron(III) oxide nanoparticles (Sigma) and 4 mM ferrozine (Sigma) prepared in mCDM-mM17. The plate was covered with aluminum foil and incubated in an anaerobic chamber (Whitley A45) at 30 °C with 150 rpm shaking. After 2 hours, 100 µL of supernatant was collected to measure the absorbance at 562 nm (Tecan Spark plate reader). The Fe^2+^ concentration was calculated using a standard curve prepared with ferrous sulfate. To test sensing plasmids, exponential-phase *L. lactis^S^*was induced with indicated concentrations of inducers in a 96-deep well plate and incubated for 15 h. Due to the difficulties in normalizing OD_600_ on the 96-deep well plate, cells were resuspended in 450 µL of mCDM-mM17, and a 13.3 µL portion was diluted to 100 µL to obtain an approximate OD_600_ of 0.2. Final concentrations of Fe^2+^ were normalized to the actual OD_600_ of *L. lactis^S^* in each well.

*E. coli^S^* induced a high background in the iron reduction assay, and thus the cell-free supernatant (CFS) was tested with *L. plantarum^R^*. *L. plantarum^R^* was resuspended in minimal mM9 to OD_600_ of 4. In a 96-deep well plate, 160 µL of *E. coli^S^* CFS was mixed with 32 µL of 20 mM iron(III) oxide nanoparticles, 32 µL of 20 mM ferrozine, and 224 µL of *L. plantarum^R^* resuspension. The plate was covered with aluminum foil and incubated under the same conditions as described above. Sensing plasmids were tested by inducing exponential-phase *E. coli^S^*with indicated concentrations of inducers in a 96-deep well plate overnight for 15 h. The final Fe^2+^ concentration was normalized to OD_600_ of overnight-grown cells in each well.

To compare the CFSs from different species, cells were grown in their outgrowth medium (**Supplementary Table 3**) before subculturing in mCDM or 1% glucose CDM (gCDM) for 15 h. *L. plantarum^R^* was subcultured in mMRS or commercial (glucose) MRS for 15 h before being washed and resuspended in mCDM or gCDM to OD_600_ of 4 to test with mCDM- or gCDM-derived CFS, respectively. CFS iron reduction assay followed the same procedure as described for *E. coli^S^*.

### Bioelectrochemical analysis of co-cultures in lab-based media

Co-cultures electroactivity and inducibility were tested in water-jacketed, two-chamber bioelectrochemical reactors (Adams & Chittenden Scientific Glass). The anodic and cathodic chambers were separated by a cation exchange membrane (CMI-7000). The anodic chamber comprised an Ag/AgCl reference electrode (CH instrument) refilled with 3M KCl saturated with silver chloride (Sigma), a 6.35-mm-thick graphite felt working electrode with a 3.2 cm radius (Alfa Aesar) threaded through a 0.5-mm radius titanium wire (Alfa Aesar), and 110 mL of medium (mCDM:mM17=20:1 for *Ll^S^-Lp^R^*co-culture or minimal mM9 for *El^S^-Lp^R^* co-culture). The cathodic chamber comprised a 0.5-mm radius titanium wire as the counter electrode and 110 mL of M9 buffer. The reactors were placed on a magnetic stir station (IKA RO 10) with the anodic chamber solution stirred at 200 rpm using a magnetic stir bar. The water jackets were connected to a heating circulator (Lauda ECO E4S), and the temperature was maintained at 30 °C. For experiments under anaerobic conditions, nitrogen (N_2_) gas was continuously sparged into the anodic chamber.

Electrochemical measurements were carried out using a VSP-300 potentiostat (BioLogic). For chronoamperometry, the working electrode was biased at +0.2 V vs. Ag/AgCl, and the current was recorded every 36 s. Once the current stabilized, 2 mL sender cells (*L. lactis^S^* or *E. coli^S^*) and 2 mL receiver cells (*L. plantarum^R^*) were injected into the anodic chamber at a final OD_600_ of 0.08 or 0.2, respectively. A 110 µL 1000-times concentrated inducer stock or blank buffer was injected 4 h post-cell injection for *L. lactis^S^* or 6 h for *E. coli^S^.* The interval time was to allow cell growth and background current stabilization. For cyclic voltammetry, the working electrode voltage was swept within the indicated range at a scan rate of 2 mV/s.

### Real-world and human-relevant sample collection and characterization

The bayou water sample was collected from the Houston area, and large solids were allowed to precipitate overnight at 4 °C before taking the supernatant for measurement. The whole milk was purchased from a local grocery store and stored at 4 °C before use. The artificial saliva was purchased from Sigma (catalog #SAE0149) and stored at 4 °C before use. All samples were not filtered for electrochemical analysis of co-cultures. The pH of each sample was measured with a pH meter (Mettler Toledo). The absorbance (A) at 509 nm was determined using a UV-Vis spectrophotometer (Agilent) with a 1 cm path-length quartz cuvette. The transmittance (T) was calculated using T = 10^(2-A). The absorbance for the milk sample was saturated, and the transmittance was near zero. To determine total organic carbon (TOC), all samples were filtered through a 0.22 µm filter (the milk sample was diluted 10^4^ times before filtering). TOC was measured using a TOC-VCSH analyzer (Shimadzu). The sample resistance was determined by the current interrupt method using a three-electrode electrochemical system. Briefly, a 10 nA current was applied to the sample, and the uncompensated resistance (R_u_) was calculated based on the ratio of the measured voltage and current (R_u_ = ΔE/ΔI).

### Human gut microbiota collection, cultivation, and metagenomic analysis

Human large intestines were obtained through the LifeGift organ donation program at the Texas Medical Center (Houston, TX, USA). All organ donors were adults without gastrointestinal disease, surgery, or trauma. Donors testing positive for hepatitis B or C, HIV, or COVID were excluded. The whole intestines were transported on ice within 1 hour of removal. Fresh large intestinal contents were resuspended in cold, sterile PBS with 20% glycerol to a final concentration of 100 mg/mL, and aliquots were stored at -80 °C until use. To cultivate large intestinal microbiotas, frozen resuspensions were thawed under anaerobic conditions. A 4 mL resuspension was inoculated into each well (6 wells total) of the minibioreactor arrays^53^ pre-filled with 15 mL of BRM3 media (**Supplementary Table 9**). Microbiota cultures were incubated for 16 h before initiating a continuous flow of fresh BRM3 media. After 5 days of continuous flow, microbiota cultures were collected from each reactor. All culturing was conducted in an anaerobic chamber with an atmosphere of 5% H₂, 5% CO₂, and 90% N₂ at 37°C.

For metagenomic analysis, cultures from the six wells of each donor’s microbiota were pooled into two groups for sequencing. Metagenomic data were analyzed using lab-developed software and ATIMA (Agile Toolkit for Incisive Microbial Analysis) developed by the Baylor College of Medicine. The remaining microbiota samples were combined for the bioelectronic sensing experiment.

### Bioelectrochemical analysis of co-cultures in real-world and human-relevant samples

Bioelectronic sensing in milk, bayou water, artificial saliva, or gut microbiota was evaluated in 5 mL-volume two-chamber bioelectrochemical reactors (Adams & Chittenden Scientific Glass). The anodic and cathodic chambers were separated by a Nafion-212 perfluorinated membrane (pre-soaked in KCl, Sigma). The working electrode was a 0.7 cm-wide, 1 cm-long, 3.18 mm-thick graphite felt (Alfa Aesar) threaded with a 0.5-mm radius titanium wire (Alfa Aesar). The reference electrode was Ag/AgCl in 3M KCl saturated with silver chloride. The counter electrode was a 0.5-mm radius titanium wire. The anodic chamber was filled with 5 mL of samples (or CDM for subsequent gut microbiota injection). The cathodic chamber was filled with 5 mL M9 buffer. The reactors were placed in an incubator at 30 °C and sparged with N_2_ gas. Current was monitored by chronoamperometry every 36 s, and the sample-derived background current was allowed to stabilize for 1 h. Exponential-phase sender and receiver cells were first mixed to create co-cultures. A 250 µL mixture of *Ll^S^-Lp^R^* co-culture in mM17 or a 500 µL mixture of *Ec^S^- Lp^R^* co-culture in minimal mM9 (2.5% mannitol) was injected into the anodic chambers for a final sender OD_600_ of 0.15 and receiver OD_600_ of 0.3. After 20 min, 25 µL 200-times concentrated inducer stock or blank buffer was injected into the anodic chambers.

To test sensing in gut microbiota, the gut microbiota samples were washed three times with N_2_-sparged 1X PBS and resuspended in N_2_-sparged 1X PBS. N_2_ sparging removed dissolved oxygen from PBS. A 100 µL resuspension was injected into the anodic chamber for a final OD_600_ of 0.25. After 1.5 h, a 250 µL mixture of *Ll^S^-Lp^R^* co-culture in mM17 was injected. Nisin or blank buffer was injected 20 min after co-culture injection.

### Screen-printed bioelectronic device manufacture

The single-chamber MFC was constructed using an Ag/AgCl-coated polyethylene Terephthalate (PET) sheet as the cathode and a porous carbon felt as the anode. The electrodes were separated by a clay proton exchange membrane (PEM). The clay PEM was made by exfoliating the thermally expanded natural vermiculite clay into atomically thin 2D flakes using a 20% aqueous HCl solution. An aqueous dispersion of these clay nanosheets was vacuum-filtered onto a cellulose nitrate membrane to fabricate a freestanding lamellar membrane. To make the anode, carbon ink (Kayaku C-250J) was screen-printed onto a PET sheet, with a 1 mm hole punched at the center. A 6 mm-diameter, 3.18 mm-thick carbon felt round was then attached to the wet carbon ink, and the ink was allowed to dry. To make the cathode, Ag/AgCl ink (Kayaku AGCL-675) was screen-printed onto a PET sheet and allowed to dry. To assemble the device, a 10 mm-diameter clay membrane was placed on top of the cathode, and an acrylic well (19 mm wide x 19 mm long x 3 mm thick) was affixed on the top of the clay membrane using glue. The carbon felt anode was then fitted into the acrylic well. The hole at the top of the anode was designed to introduce bacteria into the device.

### Screen-printed bioelectronic device signal measurements

The screen-printed devices were treated with UV/ozone for 30 min to make carbon felt hydrophilic. Exponential-phase sender and receiver cells were washed and mixed to create co-cultures. A 16.7 µL co-culture mixture was added to a 1.5 mL microcentrifuge tube containing 500 µL of testing media or samples (milk) so that the final sender OD_600_ was 1.5 and receiver OD_600_ was 3.0. For milk samples, 2.5 µL of 1 µg/mL aTc or blank ddH_2_O was added to the respective tubes before inoculating co-cultures. All tubes (three replicates for each group) were incubated at 30 °C. After 1.5-2 hours, a 60 µL portion was taken and slowly injected into the device using an insulin needle (BD). For electrochemical measurement using the potentiostat, the working electrode was connected to the carbon anode of the device, and the reference electrode was joined with the counter electrode and connected to the Ag/AgCl cathode of the device. Open circuit voltage (OCV) was recorded every 3 s. To measure short circuit current (SCA), the voltage (E) was swept from E = OCV to E = 0 vs. OCV at a scan rate of 10 mV/s. The current at E = 0 vs. OCV was taken as the SCA. For OCV measurement using the digital multimeter (Fluke 280), the positive terminal (source) was connected to the carbon anode, and the negative terminal (drain) was connected to the Ag/AgCl cathode. The OCV was measured as direct voltage (DC) in mV mode.

### Colony-forming unit counting

The sender *L. lactis^S^* or *E. coli^S^*was labeled by mCherry fluorescent protein, and the receiver *L. plantarum^R^* was labeled by superfolder green fluorescent protein (sfGFP). Co-culture samples were taken at the indicated hours and were 10-fold serially diluted from 10^1^ to 10^6^ times. A 10 µL portion from each dilution was drop-plated on a 0.5% glucose M17 plate containing 10 µg/mL erythromycin (for *Ll^S^-Lp^R^*) or 10 µg/mL chloramphenicol (for *Ec^S^-Lp^R^*) and incubated at 30 °C overnight. Colonies were counted based on fluorescence.

### Statistics

All statistical analyses were performed using the Origin Pro software (OriginLab). The p-values were determined by two-tailed unpaired Student’s t-tests, or one-way analysis of variance with Tukey’s post hoc test. Three biological replicates or more were defaulted. When two biological replicates were tested due to technical restraints, the experiments were repeated at least three times.

## Authors Contributions

Conceptualization: S.L., C.M.A.-F.

Methodology: S.L., D.Z., K.S., B.B.K.

Investigation: S.L., D.Z., K.S.

Visualization: S.L., C.M.A.-F.

Funding acquisition: S.R.S., R.A.B., C.M.A.-F.

Supervision: S.R.S., R.A.B., C.M.A.-F.

Writing – original draft: S.L., C.M.A.-F.

Writing – review editing: S.L., D.Z., K.S., B.B.K., S.R.S., R.A.B., C.M.A.-F.

## Acknowledgments

We thank Prof. Maria Marco (UC Davis) for generously sending *L. citreum*, *L. mesenteroides*, and the parental *L. lactis* strains. We are grateful to Dr. Rong Cai (Rice University) for providing guidance on the use of the 5 mL-volume two-chamber bioelectrochemical reactors; Dr. Xiao Chen and Prof. Carrie Masiello (Rice University) for assisting with the TOC measurements; Dr. Jayashree Soman (Rice University) for help with transmittance measurements; Robyn Alba (Rice University) for help with bioelectrochemical experiments; Matt Carpenter (Rice Univeristy) for providing the original H_2_O_2_ sensing plasmid; Aaron Stibelman (Rice University) for help with molecular cloning. This work was supported by the Cancer Prevention and Research Institute of Texas (award RR190063 to C.M.A.-F.) and the Army Research Office (grant W911NF-22-1-0239 to C.M.A.-F.).

## Competing interests

S.L. and C.M.A.-F. filed a provisional patent entitled “Compositions and Methods of Bioelectronic Sensing” on August 12, 2024 (No. 63/682,083), covering the design criteria of e^-^COSENS. A subsequent appendance was filed jointly with S.R.S. and K.S. on June 23, 2025 (No. 63/828,835), covering the use of the MFC device with a multimeter for electrical signal detection. S.R.S. and K.S. filed a provisional patent entitled “Nanosheet Clay Cation Exchange Membrane for Microbial Fuel Cell” on March 4, 2025 (No. 63/766,456), covering the manufacture and application of the clay membrane.

## Extended figures

**Extended Data Fig. 1.**
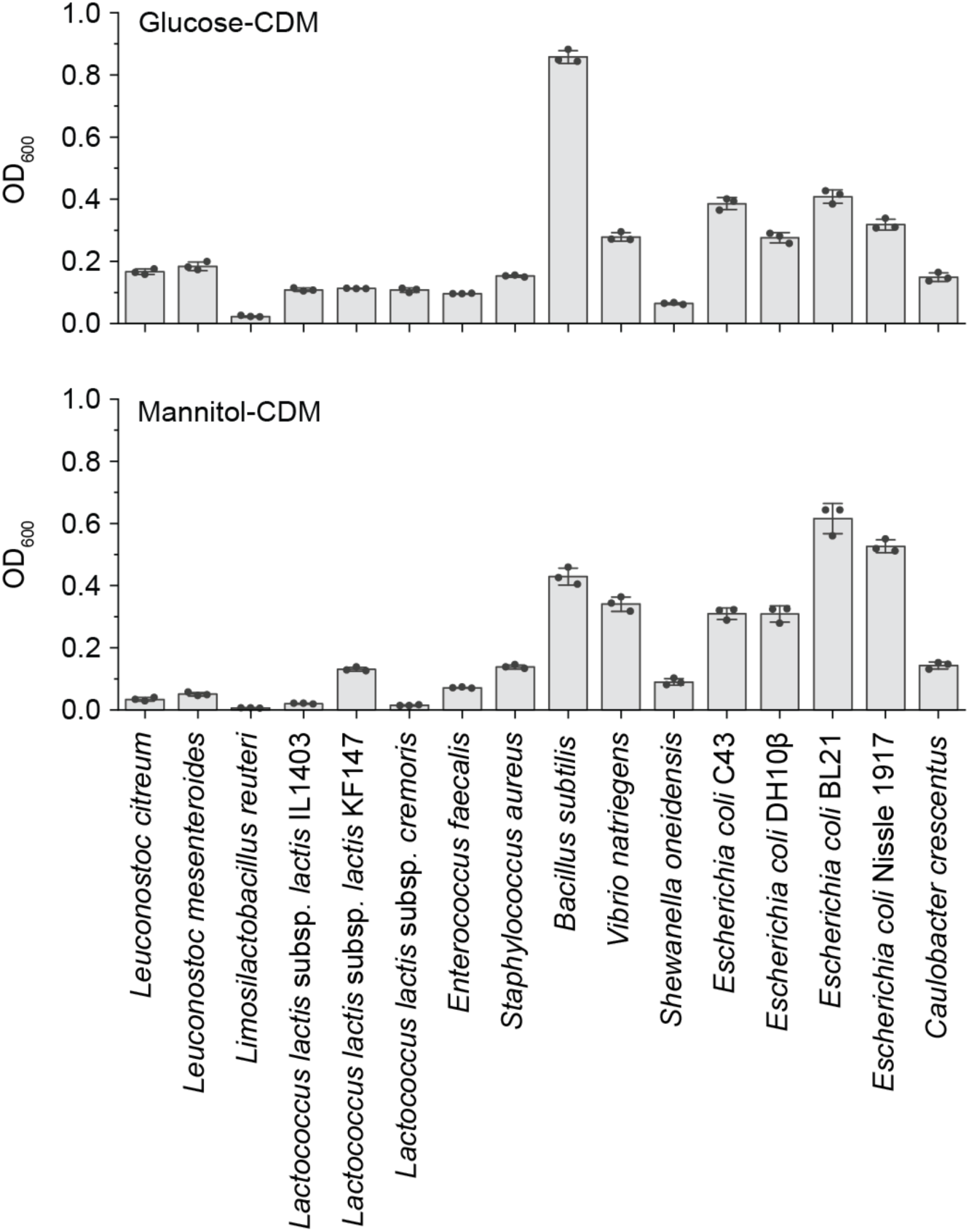
Growth of bacterial strains in chemically defined media containing 0.5% glucose or 1% mannitol. The optical density at 600 nm (OD600) of the overnight-grown bacterial cultures was determined by a plate reader. Data represent mean ± s.d. of three biological replicates.

**Extended Data Fig. 2.**
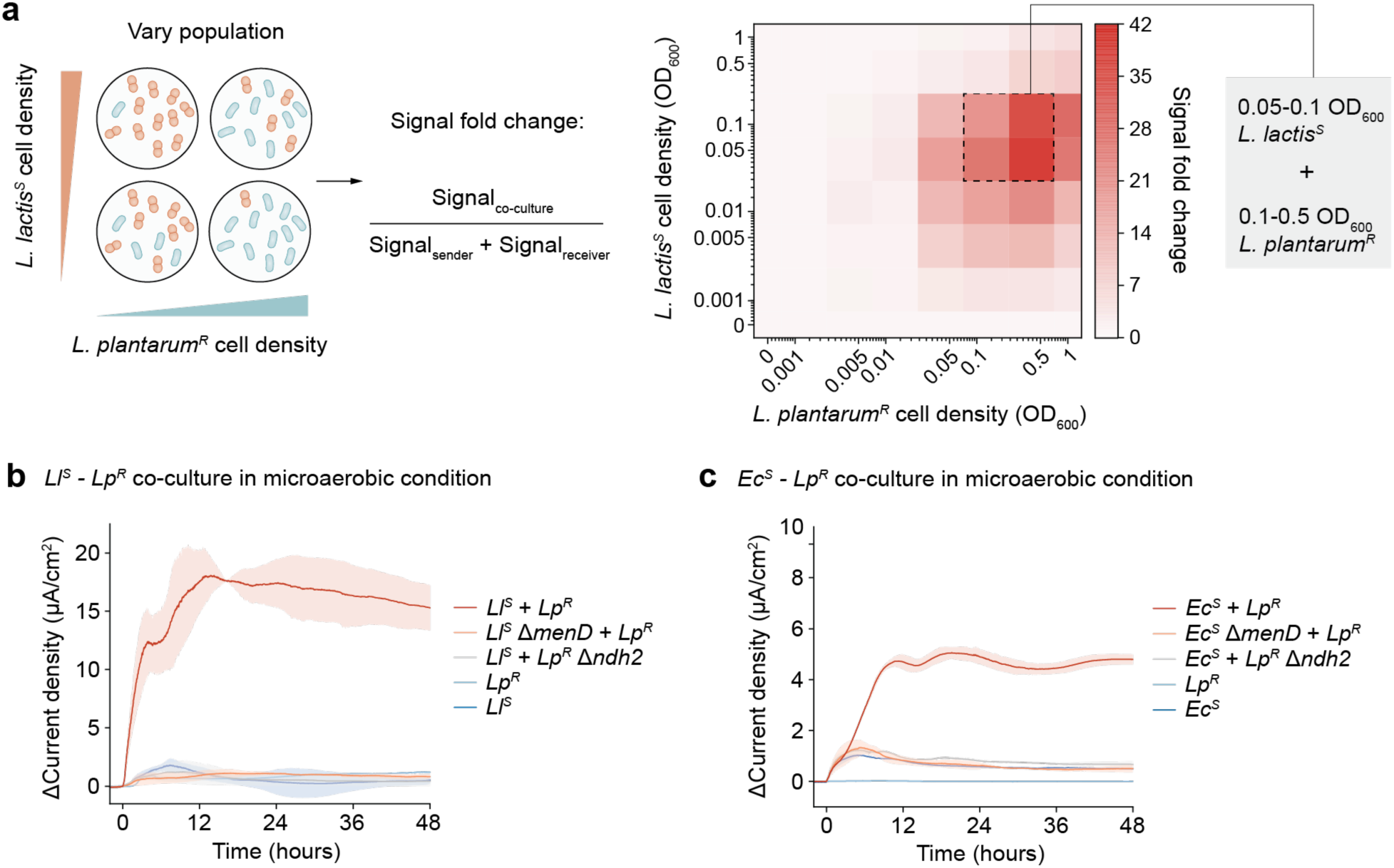
Optimization of co-culture inoculation ratio, and co-cultures produce electrical current under microaerobic conditions. **a**, Iron reduction signal fold change at various inoculation ratios of sender and receiver. The signal fold change represents the Fe^2+^ concentration reduced by the co-culture divided by the sum of the background Fe^2+^ concentration reduced by the sender or receiver monocultures. An optimal inoculation ratio is OD600 = 0.05-0.1 for the sender (*L. lactis^S^*) and OD600 = 0.1-0.5 for the receiver (*L. plantarum^R^*). Data represent the average of three biological replicates. **b,c,** Both *Ll^S^*-*Lp^R^* (**b**) and *Ec^S^*-*Lp^R^*(**c**) co-cultures produce current under microaerobic conditions (without N2 gas sparging). The current production is diminished in monocultures or when *menD* is knocked out in the sender and *ndh2* is knocked out in the receiver. Data represent mean ± s.d. of two independent biological replicates.

**Extended Data Fig. 3.**
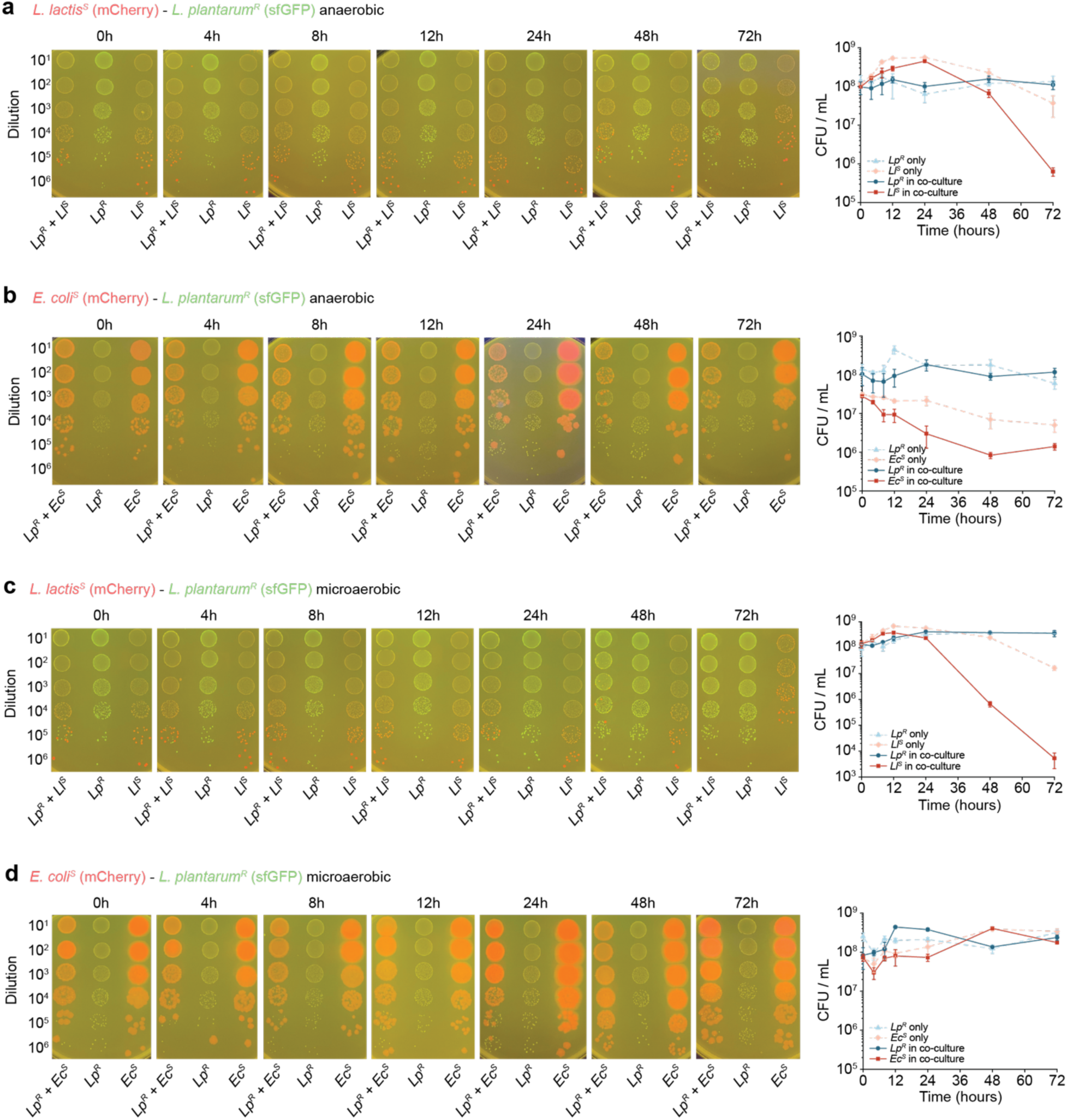
Growth curves of sender and receiver in co-cultures under anaerobic or microaerobic conditions. Growth and survival of the sender and receiver were determined by CFU counting at the indicated hours under anaerobic (**a,b**) or microaerobic (**c,d**) conditions in bioelectrochemical reactors. *L. lactis^S^* and *E. coli^S^* were labeled with mCherry, and *L. plantarum^R^* was labeled with superfolder GFP (sfGFP). Images of CFU samples are representative of three biological replicates. The quantitative results represent mean ± s.d. of three biological replicates.

**Extended Data Fig. 4.**
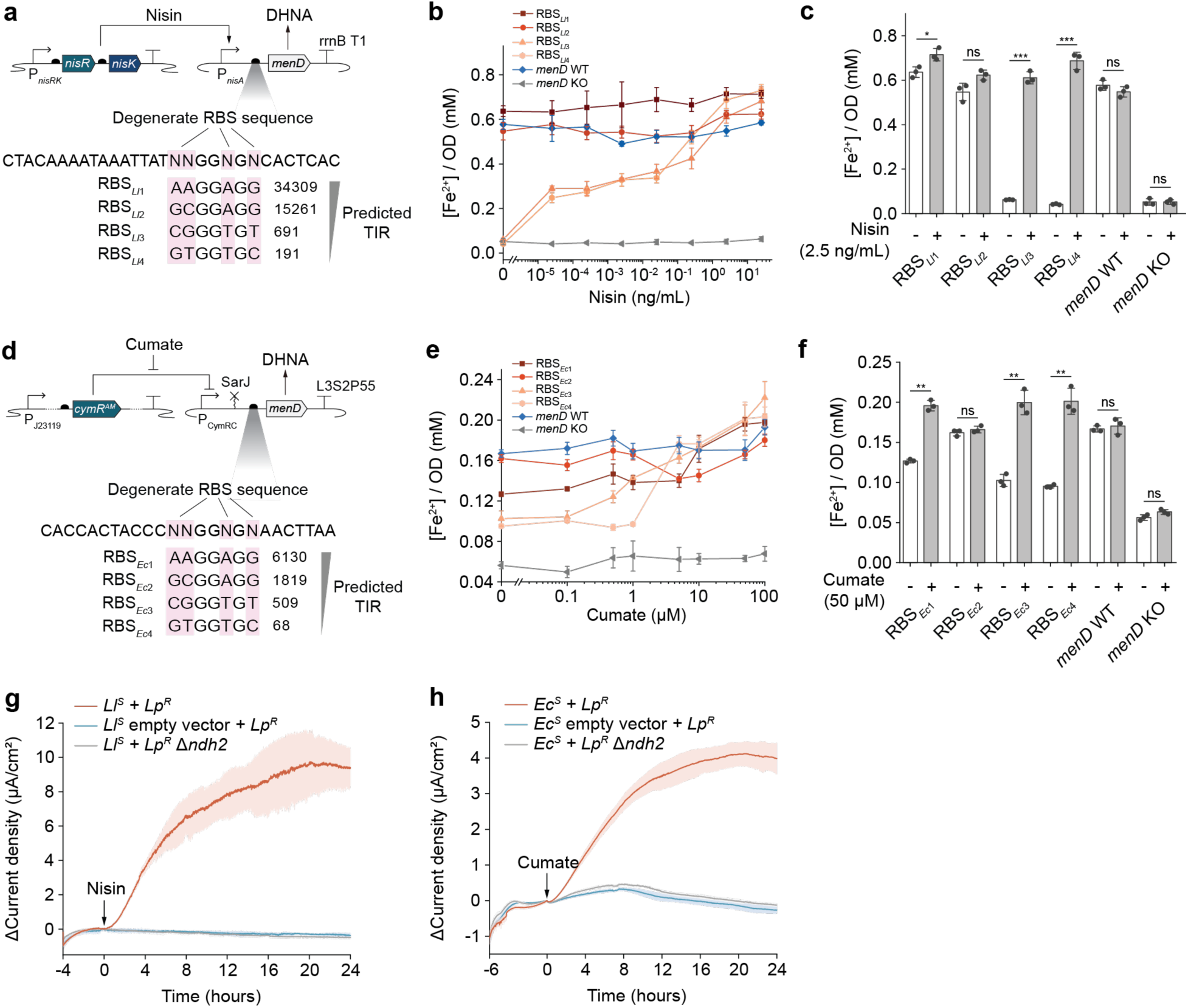
e^-^COSENS’s sensitivity can be optimized by tuning the ribosome binding site (RBS), and the analyte-induced current production depends on MenD expression and Ndh2-dependent EET. **a, d**, The translation initiation rate (TIR) of degenerate RBS (NNGGNGN) is predicted using the RBS library calculator^70,71^. Four RBSs with varied TIRs were tested to tune nisin-inducible MenD expression in *L. lactis^S^* (**a**) or cumate-inducible MenD expression in *E. coli^S^* (**d**). **b, e,** RBSs with varied TIRs result in different iron reduction fold changes upon nisin (**b**) or cumate (**e**) induction. The *menD* wild-type (genomic expression of *menD*) serves as the positive control, and the *menD* knockout serves as the negative control. Iron reduction levels were normalized to the OD600 of the sender strains. **c, f,** Quantitative analysis of iron reduction levels with or without 2.5 ng/mL nisin (**c**) or 50 µM cumate (**f**) induction. RBSs with the lowest TIR (RBS*Ll*4 and RBS*Ec*4) resulted in the highest fold change and were selected for subsequent tests. **g, h,** Current production upon nisin (2.5 ng/mL) or cumate (10 µM) induction necessitates functional co-cultures. *L. lactis^S^* or *E. coli^S^* carrying an empty vector or *L. plantarum^R^* Δ*ndh2* mutant diminished current production. All data represent mean ± s.d. of three biological replicates. P-values in **c** and **f** were determined by one-way ANOVA with Tukey’s post hoc test. *, P < 0.05; **, P < 0.01; ***, P < 0.001; ns, not significant.

**Extended Data Fig. 5.**
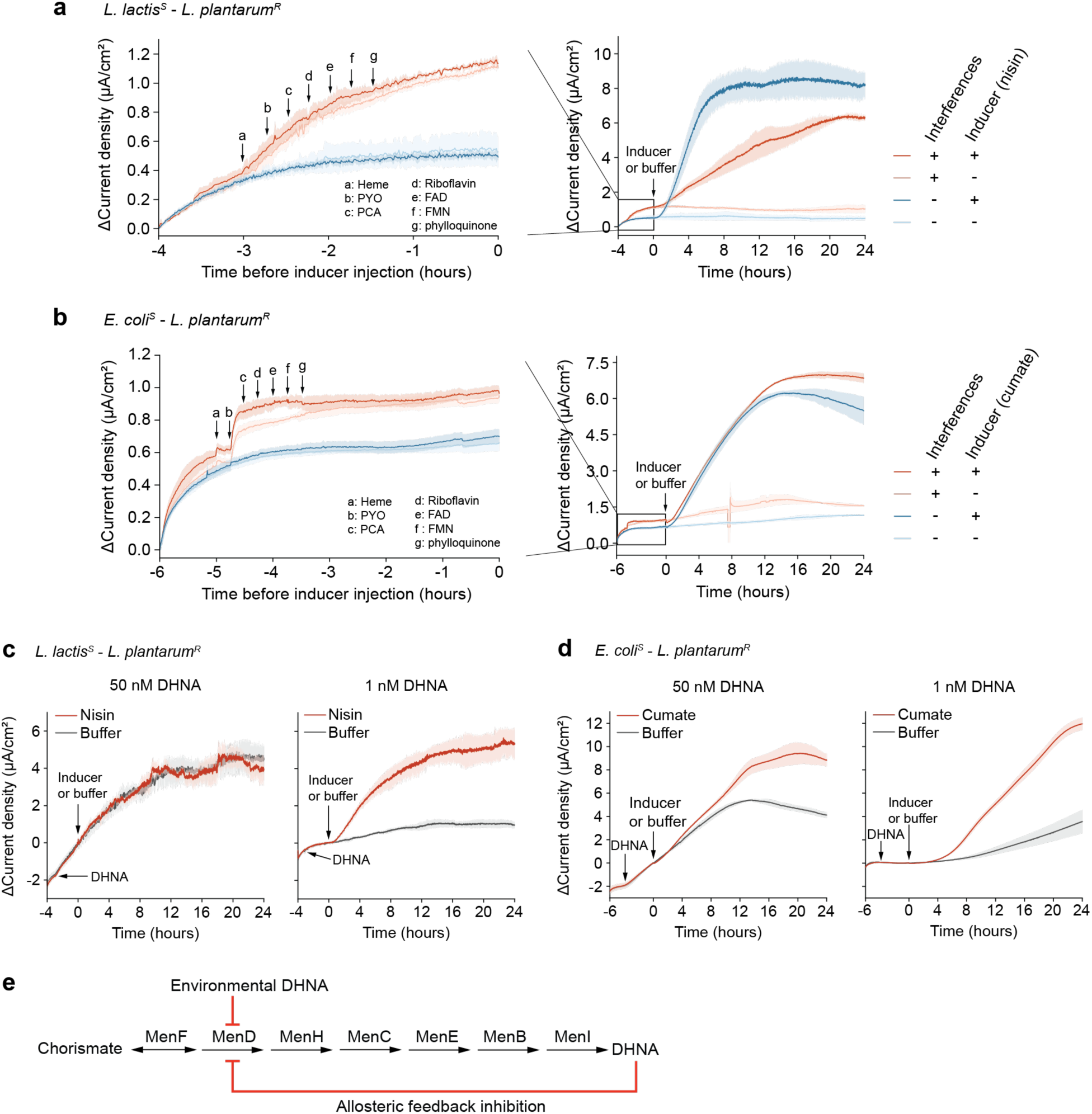
Impacts of exogenous redox-active molecules on sensing. **a,b**, Various redox-active molecules at 250 nM were sequentially added to the *Ll^S^-Lp^R^* (**a**) or *Ec^S^-Lp^R^* (**b**) co-cultures at an interval of 15 min before nisin or cumate induction. Co-cultures not exposed to redox-active molecules served as the controls. PYO, pyocyanin; PCA, phenazine-1-carboxylic acid; FAD, flavin adenine dinucleotide; FMN, flavin mononucleotide. **c, d,** Influences of exogenous DHNA at 50 or 1 nM on current production of *Ll^S^-Lp^R^* (**c**) or *Ec^S^-Lp^R^* (**d**) upon induction. DHNA of 50 nM inhibits the nisin-induced current production in *Ll^S^-Lp^R^*co-culture. **e.** Schematic showing DHNA allosterically inhibiting MenD. All data represent mean ± s.d. of three biological replicates.

**Extended Data Fig. 6.**
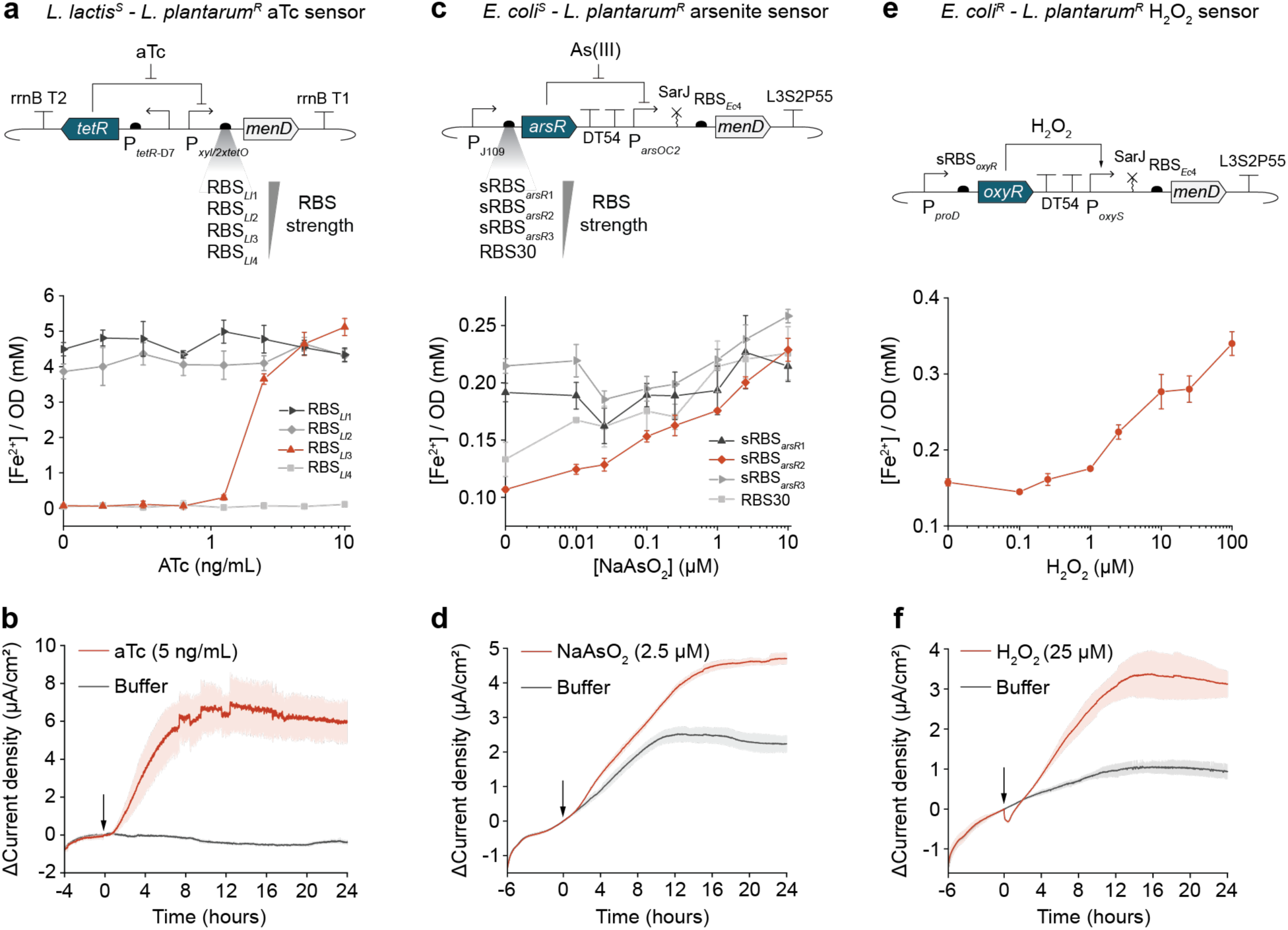
Construction, optimization, and characterization of the aTc, arsenite, and H2O2 sensors. **a**, The anhydrotetracycline (aTc) sensing genetic circuit in *L. lactis^S^* and optimization of *menD* expression by tuning RBS strength. The medium-strength RBS*Ll*3 resulted in the highest iron reduction fold change in response to varying concentrations of aTc and was selected for subsequent tests. **b,** Current production of the *Ll^S^-Lp^R^* co-culture in response to 5 ng/mL aTc. **c,** The arsenite sensing genetic circuit in *E. coli^S^* and optimization of *arsR* expression by tuning RBS strength. Synthetic RBS (sRBS) was designed using the RBS calculator^70,71^. The medium-strength sRBS*arsR*2 resulted in the highest iron reduction fold change in response to varying concentrations of arsenite (in the form of NaAsO2) and was selected for subsequent tests. **d,** Current production of the *Ec^S^-Lp^R^*co-culture in response to 2.5 µM arsenite. **e,** The hydrogen peroxide (H2O2) sensing genetic circuit in *E. coli* and iron reduction levels in response to varying concentrations of H2O2. **f.** Current production of the *Ec^S^-Lp^R^* co-culture in response to 25 µM H2O2. Arrows in **b-f** indicate inducer or buffer injection. All data represent mean ± s.d. of three biological replicates.

**Extended Data Fig. 7.**
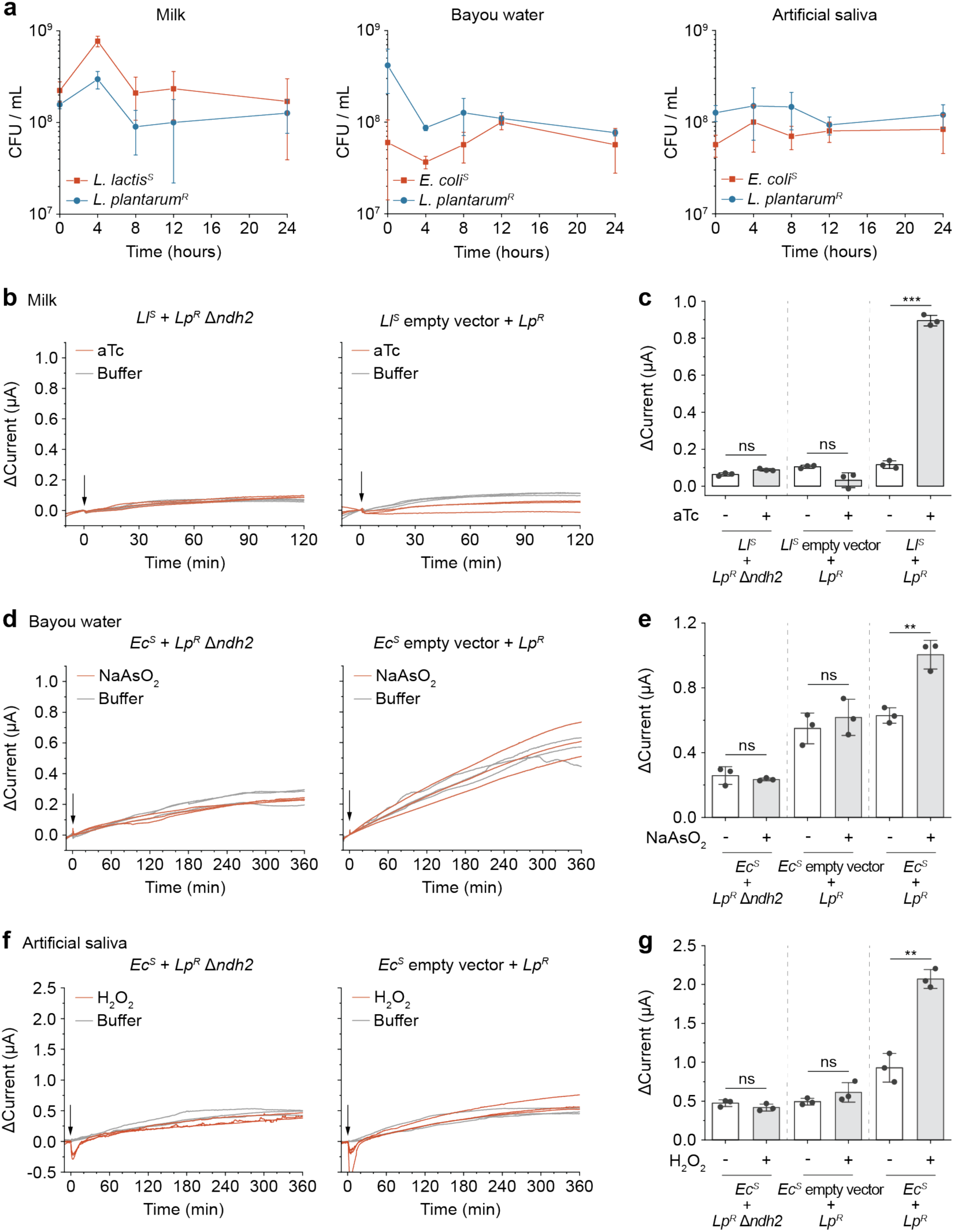
Co-culture growth analysis and current production of e^-^COSENS in milk, bayou water, and artificial saliva. **a**, CFU analysis of co-cultures at 0, 4, 8, 12, and 24 hours post-inoculation into the milk, bayou water, or artificial saliva samples. **b, d, f,** No difference in current production was observed between the induced and buffer control groups if *L. lactis* or *E. coli^S^* carried an empty vector or *ndh2* was knocked out in *L. plantarum^R^*. Arrows indicate inducer or buffer injection. **c, e, g**, Comparison of current production between deficient and functional co-cultures at 120 min (milk) or 360 min (bayou water and artificial saliva). All data represent three biological replicates or their mean ± s.d. P-values were determined by one-way ANOVA with Tukey’s post hoc test. **, P < 0.01; ***, P < 0.001; ns, not significant.

**Extended Data Fig. 8.**
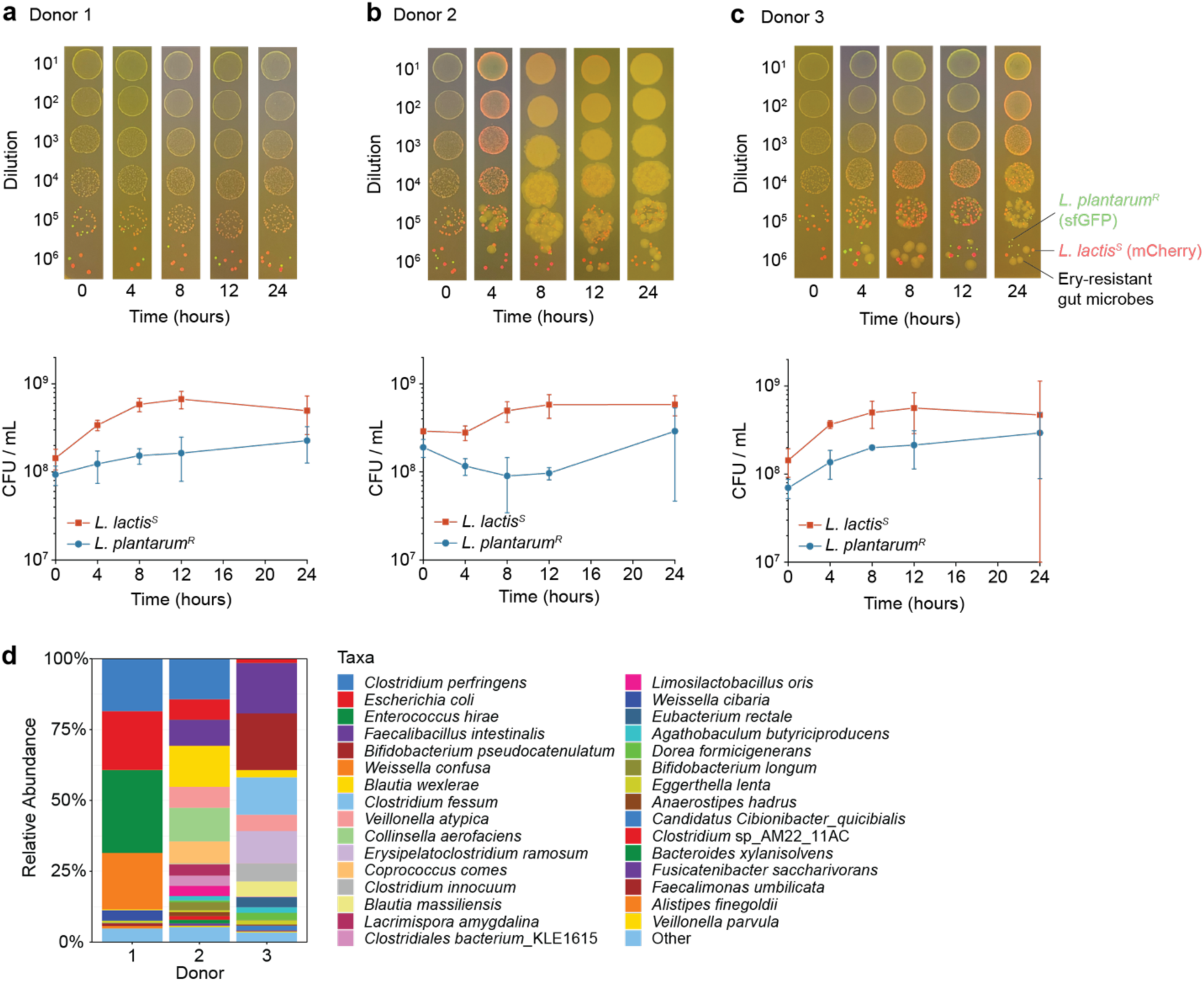
Growth analysis of *L. lactis* and *L. plantarum* co-culture in the gut microbiota and composition of gut microbiota. **a, b, c**, CFU analysis of the co-culture at 0, 4, 8, 12, and 24 hours post-inoculation into the gut microbiota from the three donors. Glucose M17 plates containing 10 µg/mL erythromycin were used for dilution plating to isolate *L. lactis^S^*and *L. plantarum^R^* from the gut microbiota. *L. plantarum^R^* was labeled with sfGFP, and *L. lactis^S^* was labeled with mCherry. The non-fluorescent colonies are erythromycin-resistant gut microbes. Images of CFU samples are representative of three biological replicates. The quantitative results represent mean ± s.d. of three biological replicates. **d,** Metagenomic analysis of the gut microbiota composition from the three human donors after four days of cultivation in the minibioreactor arrays (MBRAs).

**Extended Data Fig. 9.**
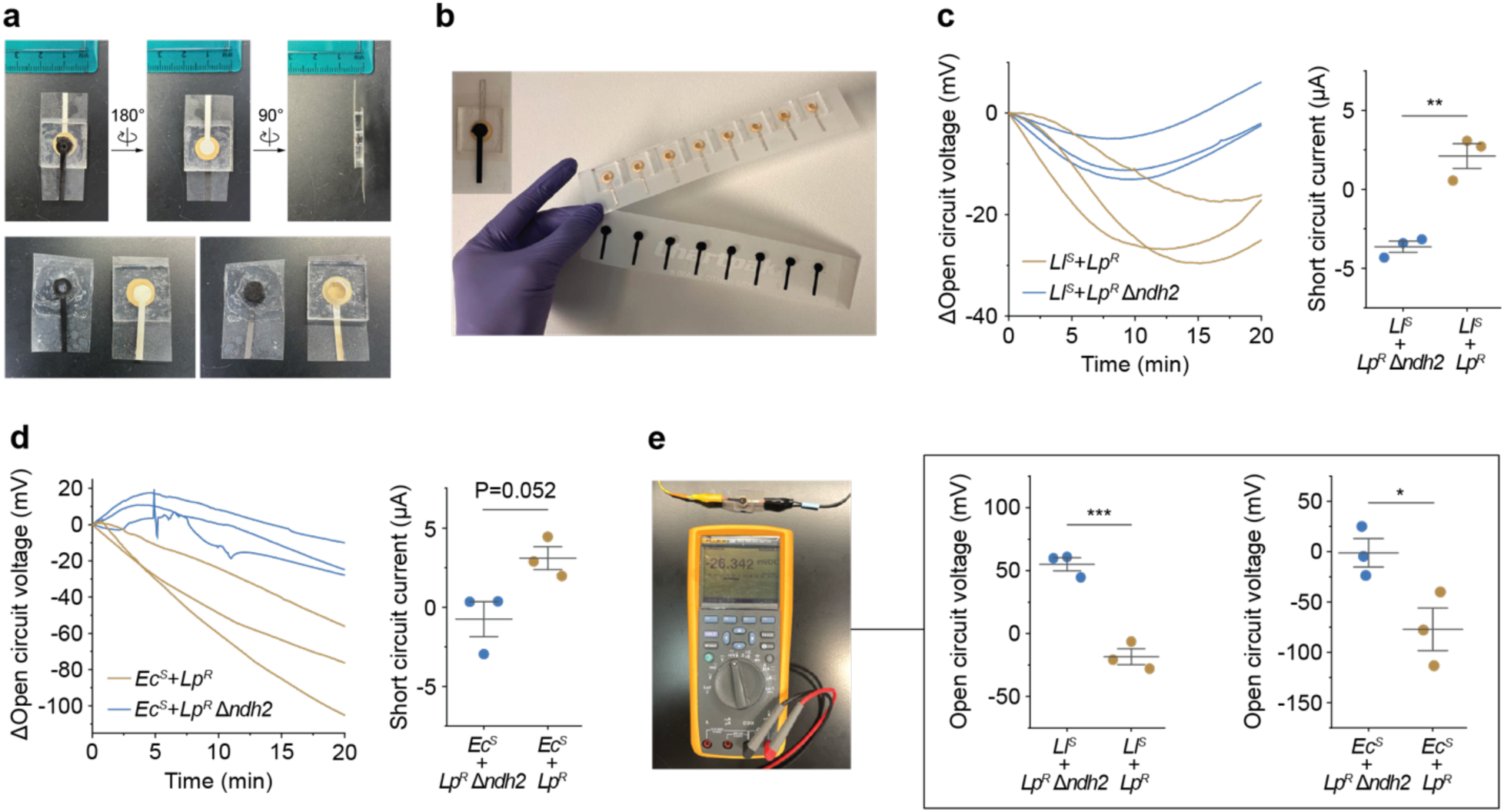
The clay-based MFC device enables signal detection from both *L. lactis-L. plantarum* and *E. coli-L. plantarum* co-cultures. **a**, Photos of the device from the front, back, and sides. **b,** Photo showing high-throughput manufacture of the clay-based MFC device. **c, d,** Measurements of the open circuit voltage (OCV) and short circuit current (SCA) for EET-capable and incapable *Ll^S^-Lp^R^* (**c**) or *Ec^S^-Lp^R^* (**d**) co-cultures**. e,** The OCV measured by the digital multimeter differentiates the EET-capable and incapable *Ll^S^-Lp^R^* or *Ec^S^-Lp^R^*co-cultures. Data represent three biological replicates or their mean ± s.e.m. P-values were determined by a two-tailed unpaired t-test. *, P < 0.05; **, P < 0.01; ***, P < 0.001.

## References

1. Fuller, R. et al. Pollution and health: a progress update. The Lancet Planetary Health 6, e535– e547 (2022).

2. Inda, M. E. & Lu, T. K. Microbes as Biosensors. Annual Review of Microbiology 74, 337–359 (2020).

3. Maresca, D. et al. Biomolecular Ultrasound and Sonogenetics. Annual Review of Chemical and Biomolecular Engineering 9, 229–252 (2018).

4. Li, S., Zuo, X., Carpenter, M. D., Verduzco, R. & Ajo-Franklin, C. M. Microbial bioelectronic sensors for environmental monitoring. Nat Rev Bioeng 3, 30–49 (2025).

5. Atkinson, J. T. et al. Real-time bioelectronic sensing of environmental contaminants. Nature 611, 548–553 (2022).

6. Karbelkar, A. A., Reynolds, E. E., Ahlmark, R. & Furst, A. L. A Microbial Electrochemical Technology to Detect and Degrade Organophosphate Pesticides. ACS Cent. Sci. 7, 1718–1727 (2021).

7. Webster, D. P. et al. An arsenic-specific biosensor with genetically engineered Shewanella oneidensis in a bioelectrochemical system. Biosensors and Bioelectronics 62, 320–324 (2014).

8. Li, S., De Groote Tavares, C., Tolar, J. G. & Ajo-Franklin, C. M. Selective bioelectronic sensing of pharmacologically relevant quinones using extracellular electron transfer in Lactiplantibacillus plantarum. Biosensors and Bioelectronics 243, 115762 (2024).

9. Gao, Y., Ryu, J., Liu, L. & Choi, S. A simple, inexpensive, and rapid method to assess antibiotic effectiveness against exoelectrogenic bacteria. Biosensors and Bioelectronics 168, 112518 (2020).

10. Cho, J. H., Gao, Y., Ryu, J. & Choi, S. Portable, Disposable, Paper-Based Microbial Fuel Cell Sensor Utilizing Freeze-Dried Bacteria for In Situ Water Quality Monitoring. ACS Omega 5, 13940–13947 (2020).

11. Chouler, J., Cruz-Izquierdo, Á., Rengaraj, S., Scott, J. L. & Di Lorenzo, M. A screen-printed paper microbial fuel cell biosensor for detection of toxic compounds in water. Biosensors and Bioelectronics 102, 49–56 (2018).

12. Pasternak, G., Greenman, J. & Ieropoulos, I. Self-powered, autonomous Biological Oxygen Demand biosensor for online water quality monitoring. Sens Actuators B Chem 244, 815–822 (2017).

13. Adekunle, A., Rickwood, C. & Tartakovsky, B. Online monitoring of heavy metal–related toxicity using flow-through and floating microbial fuel cell biosensors. Environ Monit Assess 192, 52 (2020).

14. Liu, L., Lu, Y., Zhong, W., Meng, L. & Deng, H. On-line monitoring of repeated copper pollutions using sediment microbial fuel cell based sensors in the field environment. Science of The Total Environment 748, 141544 (2020).

15. Song, N. et al. Development of a sediment microbial fuel cell-based biosensor for simultaneous online monitoring of dissolved oxygen concentrations along various depths in lake water. Science of The Total Environment 673, 272–280 (2019).

16. Mohammadifar, M. & Choi, S. A Portable and Visual Electrobiochemical Sensor for Lactate Monitoring in Sweat. in 2018 IEEE 12th International Conference on Nano/Molecular Medicine and Engineering (NANOMED) 73–77 (Waikiki Beach, HI, USA, 2018). doi:10.1109/NANOMED.2018.8641665.

17. Ueki, T., Nevin, K. P., Woodard, T. L. & Lovley, D. R. Genetic switches and related tools for controlling gene expression and electrical outputs of Geobacter sulfurreducens. Journal of Industrial Microbiology and Biotechnology 43, 1561–1575 (2016).

18. Bird, L. J. et al. Engineering Wired Life: Synthetic Biology for Electroactive Bacteria. ACS Synth. Biol. 10, 2808–2823 (2021).

19. Jensen, H. M. et al. Engineering of a synthetic electron conduit in living cells. Proc Natl Acad Sci U S A 107, 19213–19218 (2010).

20. Bird, L. J. et al. Marine Biofilm Engineered to Produce Current in Response to Small Molecules. ACS Synth. Biol. 12, 1007–1020 (2023).

21. Goldbeck, C. P. et al. Tuning promoter strengths for improved synthesis and function of electron conduits in Escherichia coli. ACS Synth Biol 2, 150–159 (2013).

22. Su, L., Fukushima, T. & Ajo-Franklin, C. M. A hybrid cyt c maturation system enhances the bioelectrical performance of engineered Escherichia coli by improving the rate-limiting step. Biosensors and Bioelectronics 165, 112312 (2020).

23. Su, L. et al. Modifying Cytochrome c Maturation Can Increase the Bioelectronic Performance of Engineered Escherichia coli. ACS Synth. Biol. 9, 115–124 (2020).

24. Light, S. H. et al. A flavin-based extracellular electron transfer mechanism in diverse Gram-positive bacteria. Nature 562, 140–144 (2018).

25. Tejedor-Sanz, S. et al. Extracellular electron transfer increases fermentation in lactic acid bacteria via a hybrid metabolism. eLife 11, e70684 (2022).

26. Seddik, H. A. et al. Lactobacillus plantarum and Its Probiotic and Food Potentialities. Probiotics Antimicrob Proteins 9, 111–122 (2017).

27. Tolar, J. G., Li, S. & Ajo-Franklin, C. M. The Differing Roles of Flavins and Quinones in Extracellular Electron Transfer in Lactiplantibacillus plantarum. Applied and Environmental Microbiology 89, e01313–22 (2022).

28. Stevens, E. T. et al. Lactiplantibacillus plantarum uses ecologically relevant, exogenous quinones for extracellular electron transfer. mBio 14, e02234–23 (2023).

29. Li, S., Zhang, J., Ajo-Franklin, C. M. & Igoshin, O. A. The growth benefits and toxicity of quinone biosynthesis are balanced by a dual regulatory mechanism and substrate limitations. mBio 16, e00887–25 (2025).

30. Kang, M.-J., Baek, K.-R., Lee, Y.-R., Kim, G.-H. & Seo, S.-O. Production of Vitamin K by Wild-Type and Engineered Microorganisms. Microorganisms 10, 554 (2022).

31. Liu, Y., van Bennekom, E. O., Zhang, Y., Abee, T. & Smid, E. J. Long-chain vitamin K2 production in Lactococcus lactis is influenced by temperature, carbon source, aeration and mode of energy metabolism. Microbial Cell Factories 18, 129 (2019).

32. Song, A. A.-L., In, L. L. A., Lim, S. H. E. & Rahim, R. A. A review on Lactococcus lactis: from food to factory. Microbial Cell Factories 16, 55 (2017).

33. Meyer, A. J., Segall-Shapiro, T. H., Glassey, E., Zhang, J. & Voigt, C. A. Escherichia coli “Marionette” strains with 12 highly optimized small-molecule sensors. Nat Chem Biol 15, 196–204 (2019).

34. Andrews, S. C., Robinson, A. K. & Rodríguez-Quiñones, F. Bacterial iron homeostasis. FEMS Microbiology Reviews 27, 215–237 (2003).

35. Alba, R. A. C., Li, S., Kundu, B. B., Ajo-Franklin, C. M. & Cai, R. Characterizing Mediated Extracellular Electron Transfer in Lactic Acid Bacteria with a Three-Electrode, Two-Chamber Bioelectrochemical System. Journal of Visualized Experiments e67204 (2024) doi:10.3791/67204.

36. de Ruyter, P. G., Kuipers, O. P. & de Vos, W. M. Controlled gene expression systems for Lactococcus lactis with the food-grade inducer nisin. Appl Environ Microbiol 62, 3662–3667 (1996).

37. VanArsdale, E. et al. A Coculture Based Tyrosine-Tyrosinase Electrochemical Gene Circuit for Connecting Cellular Communication with Electronic Networks. ACS Synth. Biol. 9, 1117–1128 (2020).

38. Tschirhart, T. et al. Electrochemical Measurement of the β-Galactosidase Reporter from Live Cells: A Comparison to the Miller Assay. ACS Synth. Biol. 5, 28–35 (2016).

39. Zhou, T. et al. A copper-specific microbial fuel cell biosensor based on riboflavin biosynthesis of engineered Escherichia coli. Biotechnology and Bioengineering 118, 210–222 (2021).

40. Khan, A. et al. A novel biosensor for zinc detection based on microbial fuel cell system. Biosensors and Bioelectronics 147, 111763 (2020).

41. Bashiri, G. et al. Allosteric regulation of menaquinone (vitamin K2) biosynthesis in the human pathogen Mycobacterium tuberculosis. J Biol Chem 295, 3759–3770 (2020).

42. Stanborough, T. et al. Allosteric inhibition of Staphylococcus aureus MenD by 1,4-dihydroxy naphthoic acid: a feedback inhibition mechanism of the menaquinone biosynthesis pathway. Philosophical Transactions of the Royal Society B: Biological Sciences 378, 20220035 (2023).

43. Markakiou, S., Neves, A. R., Zeidan, A. A. & Gaspar, P. Development of a Tetracycline-Inducible System for Conditional Gene Expression in Lactococcus lactis and Streptococcus thermophilus. Microbiology Spectrum 11, e00668–23 (2023).

44. Chen, S.-Y., Zhang, Y., Li, R., Wang, B. & Ye, B.-C. De Novo Design of the ArsR Regulated Pars Promoter Enables a Highly Sensitive Whole-Cell Biosensor for Arsenic Contamination. Anal. Chem. 94, 7210–7218 (2022).

45. Rubens, J. R., Selvaggio, G. & Lu, T. K. Synthetic mixed-signal computation in living cells. Nat Commun 7, 11658 (2016).

46. Mays, Z. J. & Nair, N. U. Synthetic biology in probiotic lactic acid bacteria: At the frontier of living therapeutics. Current Opinion in Biotechnology 53, 224–231 (2018).

47. Kotula, J. W. et al. Programmable bacteria detect and record an environmental signal in the mammalian gut. Proc. Natl. Acad. Sci. U.S.A. 111, 4838–4843 (2014).

48. Mimee, M. et al. An ingestible bacterial-electronic system to monitor gastrointestinal health. Science 360, 915–918 (2018).

49. Inda-Webb, M. E. et al. Sub-1.4 cm3 capsule for detecting labile inflammatory biomarkers in situ. Nature 620, 386–392 (2023).

50. J Pytko-Polonczyk, J., Jakubik, A., Przeklasa-Bierowiec, A. & Muszynska, B. Artificial saliva and its use in biological experiments. J Physiol Pharmacol 68, 807–813 (2017).

51. Martino, M. E. et al. Nomadic lifestyle of *Lactobacillus plantarum* revealed by comparative genomics of 54 strains isolated from different habitats. Environmental Microbiology 18, 4974–4989 (2016).

52. De Filippis, F., Pasolli, E. & Ercolini, D. The food-gut axis: lactic acid bacteria and their link to food, the gut microbiome and human health. FEMS Microbiology Reviews 44, 454 (2020).

53. Auchtung, J. M., Robinson, C. D. & Britton, R. A. Cultivation of stable, reproducible microbial communities from different fecal donors using minibioreactor arrays (MBRAs). Microbiome 3, 42 (2015).

54. Bora, B. R., Nath, N., Dey, M., Saha, K. & Raidongia, K. Assembly of Natural Clay Minerals as Highly Robust Evaporation-Driven Power Generator. ACS Appl. Energy Mater. 7, 6507–6514 (2024).

55. Del Valle, I. et al. Translating New Synthetic Biology Advances for Biosensing Into the Earth and Environmental Sciences. Front. Microbiol. 11, 618373 (2021).

56. Park, Y., Espah Borujeni, A., Gorochowski, T. E., Shin, J. & Voigt, C. A. Precision design of stable genetic circuits carried in highly-insulated E. coli genomic landing pads. Molecular Systems Biology 16, e9584 (2020).

57. Kok, J. et al. The Evolution of gene regulation research in Lactococcus lactis. FEMS Microbiology Reviews 41, S220–S243 (2017).

58. Tschirhart, T. et al. Synthetic Biology Tools for the Fast-Growing Marine Bacterium *Vibrio natriegens*. ACS Synth. Biol. 8, 2069–2079 (2019).

59. Liu, Y., Liu, L., Li, J., Du, G. & Chen, J. Synthetic Biology Toolbox and Chassis Development in Bacillus subtilis. Trends in Biotechnology 37, 548–562 (2019).

60. Siezen, R. J. et al. Genome-scale diversity and niche adaptation analysis of Lactococcus lactis by comparative genome hybridization using multi-strain arrays. Microb Biotechnol 4, 383–402 (2011).

61. Hoff, J., et al. *Vibrio natriegens* : an ultrafast-growing marine bacterium as emerging synthetic biology chassis. Environmental Microbiology 22, 4394–4408 (2020).

62. Su, Y., Liu, C., Fang, H. & Zhang, D. Bacillus subtilis: a universal cell factory for industry, agriculture, biomaterials and medicine. Microbial Cell Factories 19, 173 (2020).

63. Zuo, X. et al. Quinone-Grafted Chitosan Polymers Enhance Microbial Extracellular Electron Transfer for Living Bioelectronic Devices. Preprint at 10.26434/chemrxiv-2025-ss6fz (2025).

64. Qin, X. & Taber, H. W. Transcriptional regulation of the Bacillus subtilis menp1 promoter. Journal of Bacteriology 178, 705–713 (1996).

65. Grattieri, M. & Minteer, S. D. Self-Powered Biosensors. ACS Sens. 3, 44–53 (2018).

66. Leloup, L., Ehrlich, S. D., Zagorec, M. & Morel-Deville, F. Single-crossover integration in the Lactobacillus sake chromosome and insertional inactivation of the ptsI and lacL genes. Appl Environ Microbiol 63, 2117–2123 (1997).

67. Li, Q. et al. A modified pCas/pTargetF system for CRISPR-Cas9-assisted genome editing in Escherichia coli. Acta Biochimica et Biophysica Sinica 53, 620–627 (2021).

68. Kuipers, O. P., de Ruyter, P. G. G. A., Kleerebezem, M. & de Vos, W. M. Quorum sensing-controlled gene expression in lactic acid bacteria. Journal of Biotechnology 64, 15–21 (1998).

69. Engler, C., Kandzia, R. & Marillonnet, S. A One Pot, One Step, Precision Cloning Method with High Throughput Capability. PLoS ONE 3, e3647 (2008).

70. Salis, H. M., Mirsky, E. A. & Voigt, C. A. Automated design of synthetic ribosome binding sites to control protein expression. Nat Biotechnol 27, 946–950 (2009).

71. Reis, A. C. & Salis, H. M. An Automated Model Test System for Systematic Development and Improvement of Gene Expression Models. ACS Synth. Biol. 9, 3145–3156 (2020).

72. Sørvig, E., Mathiesen, G., Naterstad, K., Eijsink, V. G. H. & Axelsson, L. High-level, inducible gene expression in Lactobacillus sakei and Lactobacillus plantarum using versatile expression vectors. *Microbiology (Reading*, Engl.) 151, 2439–2449 (2005).

73. Bansal, P. et al. Rhea, the reaction knowledgebase in 2022. Nucleic Acids Res 50, D693–D700 (2022).

74. Hunter, S. et al. InterPro: the integrative protein signature database. Nucleic Acids Research 37, D211 (2008).

75. Li, W. et al. RefSeq: expanding the Prokaryotic Genome Annotation Pipeline reach with protein family model curation. Nucleic Acids Res 49, D1020–D1028 (2021).

76. Parks, D. H. et al. A standardized bacterial taxonomy based on genome phylogeny substantially revises the tree of life. Nat Biotechnol 36, 996–1004 (2018).

