## Supplementary Information for "Synthetic microbial co-cultures for modular bioelectronic sensing in diverse environments"

This PDF includes:

### Supplementary Figures

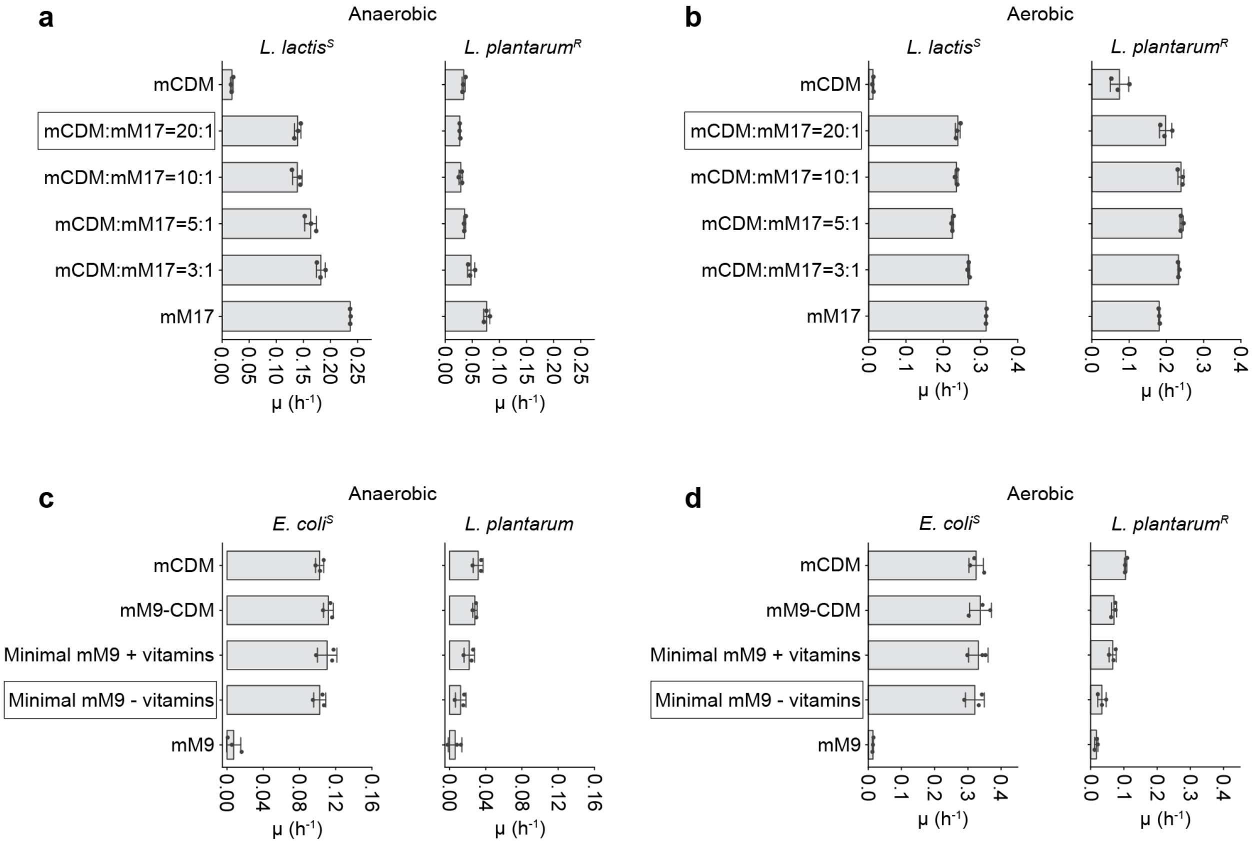

**Supplementary Fig. 1 | Optimization of co-culture media.** The medium was optimized to 1) support the growth of the sender strains (*L. lactis*$^S$ or *E. coli*$^S$) for efficient transcriptional regulation in sensing; 2) allow the receiver strain (*L. plantarum*$^R$) to maintain metabolic activity without requiring growth; 3) keep the medium composition as simple as possible to minimize the electrochemical background. **a, b,** Specific growth rate (μ) of *L. lactis*$^S$ or *L. plantarum*$^R$ in blended mCDM and mM17 media under anaerobic (**a**) or aerobic (**b**) conditions. A medium composed of mCDM and mM17 in a ratio of 20:1 was selected for co-culture, as it provides the simplest composition to support *L. lactis*$^S$ growth while maintaining the activity of *L. plantarum*$^R$*.* **c, d,** Specific growth rate (μ) of *E. coli*$^S$ or *L. plantarum*$^R$ in M9-derived media under anaerobic (**a**) or aerobic (**b**) conditions. mM9-CDM was made by substituting the MOPS buffer and salts in mCDM (**Supplementary Table 6**) with M9 salts. Minimal mM9 (with vitamins) was made by supplementing 1X Wolfe's vitamin solution to the minimal mM9. mM9 was made by adding 1% mannitol to the M9 salt solution. The minimal mM9 (without vitamins) was selected for co-culture, as it offers the simplest composition to support *E. coli*$^S$ growth while maintaining the activity of *L. plantarum*$^R$. All data represent mean ± s.d. of three biological replicates.

### Supplementary Tables

**Supplementary Table 1 | List of bacteria that possess a DHNA biosynthesis pathway**
(Supplementary Table 1 is provided as a separate file)

**Supplementary Table 2 | List of Firmicutes with an EET locus but possess an incomplete or missing DHNA biosynthesis pathway**
(Supplementary Table 2 is provided as a separate file)

**Supplementary Table 3 | List of strains**

| **Strains for assembling electroactive co-cultures**<br>(Note: sensing strains are constructed by transforming the respective plasmid listed in Supplementary Table 4 into the sender Δ*menD* strains) | | |
|---|---|---|
| **Strain** | **Description** | **Ref.** |
| *Lactiplantibacillus plantarum* NCIMB8826 Δ*dmkA* Δ*ndh1* | Receiver *L. plantarum*$^R$ | [1] |
| *Lactiplantibacillus plantarum* NCIMB8826 Δ*dmkA* Δ*ndh1* Δ*ndh2* | Receiver *L. plantarum*$^R$ deficient in EET (Δ*ndh2*) | [1] |
| *Lactococcus lactis* subsp. *lactis* KF147 Δ*noxAB* Δ*menA* | Sender *L. lactis*$^S$ | This work |
| *Lactococcus lactis* subsp. *lactis* KF147 Δ*noxAB* Δ*menA* Δ*menD* | Sender *L. lactis*$^S$ deficient in DHNA biosynthesis (Δ*menD*) | This work |
| *Escherichia coli* “Marionette” BL21 Δ*menA* | Sender *E. coli*$^S$ | “Marionette” BL21[2]<br>Mutant: This work |
| *Escherichia coli* “Marionette” BL21 Δ*menA* Δ*menD* | Sender *E. coli*$^S$ deficient in DHNA biosynthesis (Δ*menD*) | “Marionette” BL21[2]<br>Mutant: This work |
| **Strains for testing cell-free supernatant** | | |
| **Strain** | **Outgrowth medium** | |
| *Leuconostoc citreum* NRRL-B742 | MRS | |
| *Leuconostoc mesenteroides* ATCC8293 | MRS | |
| *Limosilactobacillus reuteri* subsp. *reuteri* ATCC PTA 6475 | MRS | |
| *Lactococcus lactis* subsp. *lactis* IL1403 | glucose-M17 (gM-17) | |
| *Lactococcus lactis* subsp. *lactis* KF147 | glucose-M17 (gM-17) | |
| *Lactococcus lactis* subsp. *cremoris* MG1363 | glucose-M17 (gM-17) | |
| *Enterococcus faecalis* OG1RF | Tryptic soy broth (TSB) | |
| *Staphylococcus aureus* MRSA 371 | Tryptic soy broth (TSB) | |
| *Bacillus subtilis* 168 | LB | |
| *Vibrio natriegens* ATCC 14048 | LB v2 | |
| *Shewanella oneidensis* MR-1 | LB | |
| *Escherichia coli* C43 | LB | |
| *Escherichia coli* DH10β | LB | |
| *Escherichia coli* BL21 | LB | |
| *Escherichia coli* Nissle 1917 | LB | |
| *Caulobacter crescentus* NA1000 | PYE | |

**Supplementary Table 4 | List of plasmids**

| **Plasmid** | **Insertion details** | **Function** |
|---|---|---|
| **Plasmids for *L. lactis*$^{S}$** (backbone: ColE1 ori_*ampR*_AMβ1 ori_*ermR*) | | |
| pSL145 | $P_{nisA}$_RBS$_{Ll1}$_*menD* | Nisin sensing with RBS$_{Ll1}$ |
| pSL146 | $P_{nisA}$_RBS$_{Ll2}$_*menD* | Nisin sensing with RBS$_{Ll2}$ |
| pSL147 | $P_{nisA}$_RBS$_{Ll3}$_*menD* | Nisin sensing with RBS$_{Ll3}$ |
| pSL137 | $P_{nisA}$_RBS$_{Ll4}$_*menD* | Nisin sensing with RBS$_{Ll4}$ |
| pSL187 | *tetR*_$P_{tetR\text{-}D7}$_$P_{xyl/2xtetO}$_RBS$_{Ll1}$_*menD* | ATc sensing with RBS$_{Ll1}$ |
| pSL186 | *tetR*_$P_{tetR\text{-}D7}$_$P_{xyl/2xtetO}$_RBS$_{Ll2}$_*menD* | ATc sensing with RBS$_{Ll2}$ |
| pSL185 | *tetR*_$P_{tetR\text{-}D7}$_$P_{xyl/2xtetO}$_RBS$_{Ll3}$_*menD* | ATc sensing with RBS$_{Ll3}$ |
| pSL181 | *tetR*_$P_{tetR\text{-}D7}$_$P_{xyl/2xtetO}$_RBS$_{Ll4}$_*menD* | ATc sensing with RBS$_{Ll4}$ |
| pSL141 | $P_{23}$_*mCherry* | Label *L. lactis* with mCherry |
| pSL076 | / | Empty vector for *L. lactis* |
| **Plasmids for *E. coli*$^{S}$** (backbone: p15A ori_*cmR*) | | |
| pSL160 | $P_{CymRC}$_SarJ_RBS$_{Ec1}$_*menD* | Cumate sensing with RBS$_{Ec1}$ |
| pSL161 | $P_{CymRC}$_SarJ_RBS$_{Ec2}$_*menD* | Cumate sensing with RBS$_{Ec2}$ |
| pSL162 | $P_{CymRC}$_SarJ_RBS$_{Ec3}$_*menD* | Cumate sensing with RBS$_{Ec3}$ |
| pSL163 | $P_{CymRC}$_SarJ_RBS$_{Ec4}$_*menD* | Cumate sensing with RBS$_{Ec4}$ |
| pSL177 | $P_{J109}$_RBS30_*arsR*_$P_{arsOC2}$_SarJ_RBS$_{Ec4}$_*menD* | Arsenite sensing with RBS30 |
| pSL178 | $P_{J109}$_sRBS$_{arsR1}$_*arsR*_$P_{arsOC2}$_SarJ_RBS$_{Ec4}$_*menD* | Arsenite sensing with sRBS$_{arsR1}$ |
| pSL179 | $P_{J109}$_sRBS$_{arsR2}$_*arsR*_$P_{arsOC2}$_SarJ_RBS$_{Ec4}$_*menD* | Arsenite sensing with sRBS$_{arsR2}$ |
| pSL180 | $P_{J109}$_sRBS$_{arsR3}$_*arsR*_$P_{arsOC2}$_SarJ_RBS$_{Ec4}$_*menD* | Arsenite sensing with sRBS$_{arsR3}$ |
| pSL170 | $P_{proD}$_sRBS$_{oxyR}$_*oxyR*_$P_{oxyS}$_SarJ_RBS$_{Ec4}$_*menD* | $H_2O_2$ sensing plasmid |
| pSL157 | $P_{J23101}$_RiboJ00_RBS34_*mCherry2* | Label *E. coli* with mCherry |
| pSL168 | / | Empty vector for *E. coli* |
| **Plasmids for *L. plantarum*$^{R}$** | | |
| pCD013 | pUC(pGEM) ori_256rep_*ermR*_*ampR*_$P_{tuf}$_*sfgfp* | Label *L. plantarum* with sfGFP (Ery-resistant) |
| pCD002 | pUC(pGEM) ori_256rep_*ermR*_*ampR* | Erythromycin-resistant empty vector for *L. plantarum* |
| pSL144 | pUC(pGEM) ori_256rep_*cat*_*ampR*_$P_{23}$_*sfgfp* | Label *L. plantarum* with sfGFP (Chl-resistant) |
| pSL143 | pUC(pGEM) ori_256rep_*cat*_*ampR* | Chloramphenicol-resistant empty vector for *L. plantarum* |
| **Plasmids for genome modification** | | |
| pRV300 | Homologous arms for *noxAB*, *menA*, or *menD* knockout in *L. lactis* were inserted into the NotI-EcoRI digested sites | Suicide plasmid for double-crossover homologous recombination[3] |
| pEcCas (Addgene #73227) | / | Expression of Cas and λ-Red recombinase[4] |
| pTargetF (Addgene #62226) | sgRNA was replaced to target *menA* or *menD* in *E. coli* | Expression of sgRNA[5] |

### Supplementary Table 5 | Sequence of genetic parts

| Part name | Part class | Sequence (5'-3') |
|---|---|---|
| **Parts for *L. lactis*[S]** | | |
| $P_{nisA}$ | Promoter | agtcttataactatactgacaatagaaacattaacaaatctaaaacagtcttaattctatcttgagaaagtattggtaataatattattgtcgataacgcgagcataataaacggctctgattaaattctgaagtttgttagatacaatgatttcgttcgaaggaa |
| $P_{xyl/2xtetO}$ | Promoter | ctcgagttcatgaaaaactaaaaaaaatattgacactctatcattgatagagtataattaaaataagctctctatcattgatagagtatgatggtac |
| $P_{tetR\text{-}D7}$ | Promoter + RBS | gggatccaaataaaaaactagtttgacaaataactcatcaatgatataatgtcaacaaaacaaaggaattaatg |
| $P_{23}$ | Promoter + RBS | cgaaaagccctgacaacccttgttcctaaaaaggaataagcgttcggtcagtaaataatagaaataaaaaatcagacctaagactgatgacaaaaagagaaattttgataaaatagtcttagaattaaattaaaaagggaggccaaatata |
| $RBS_{Ll1}$ | RBS | ctacaaaataaattatAAGGAGGcactcac |
| $RBS_{Ll2}$ | RBS | ctacaaaataaattatGCGGAGGcactcac |
| $RBS_{Ll3}$ | RBS | ctacaaaataaattatCGGGTGTcactcac |
| $RBS_{Ll4}$ | RBS | ctacaaaataaattatGTGGTGCcactcac |
| *menD* (*L. lactis* KF147) | Gene | atgaccaatgaatatttagctccctttgttgatgaacttttcaatttaggggtacgtgaagctgtttttagtccaggttcacgttcaacggctttagccatgctctttgaggaatacaaaaaatacgatacttatgtcaacattgatgaacgttcggctgcttttttgctctgggaattgctaaagcgaaaaaacgtccttgttgttttagtttgtacttcaggttctgctgcggctcatcattttccagctgtccttgaagcaaaaatgagtcgagttccccttattattttaacggcggacagaccggctgagcttcaattcgttggagctcctcaaacagtcgaccagaccagatttttggaaattttgttaatcattttgaaaatcttgaagctccacatattcaaacccagtctcagacagaaaacttttggacctatccacgtaaagtcgcacaacgagctgtcttatcagctatttcccctctttcgggtcctgtgcaaatcaatgttcccttgcgtgatcctttagttccagaattaaaaagtgaaaattatgaaaaaggacgttctaaacatgcctttaaattctataagggacaagcacaagtaatacttccatttgatgaaacattgctttctggaaaaacattaattttagccggtgccaattttgacaaagattattctgaatcacttttaaaacttgcggaacaattaaaagctccaattttagcggatccactttctaatctgcgtaatcataatcatcctttggtcatagactcttatgatgcttttttagcaaataatgatttaaaaattcaattaaaagcggatgctattctgctttttggacaaatgcctgtttctaaacgtttgcaacaattcgttgctttcaataacgaagctgaatttattcaagttgaccctgctcttgattatcgcaaccctagtttgacaacgacaacgatggttcagtctgatgttctcccttttgcttcttcaattaaaaaagtaaattctgatagttcttatttggaaaaatggcaaaatgctcaagaaaaaatgcgacaacttcttgaaaaagtgacttatgaagaaagtccttttgaaggacgttatgtccaagaactacaaaaacatttggaagcactggatgctcaacttcttgtttcaaatagtatggagatcagagatattgattattggtggaaaaaagcggatgcaaaagttagaattctaggaaatcgtggtgtgaatggcattgacgggactgaatcaactgcactcggaattgctacaacgggtaaaccgacggttttattgactggggatttgtcaatgctacacgatttaaatggtttaattgttggtaaaacacatgaacttaatctgactattgtgctatttaataatgacggtggtgggatttttcatcatttagctcaaaaaggcgtaccgaattttgattatctttttcaactccacatggtttaaattttgctggactggctgagttgactggtttagattatcatttggtcagtggctatgctgattttggtcagcaatttgaagcttctcttcatcaatctggcattcatcttttggaaattaaaacggataaagatttaagtcttgctttacatcaaaaatatacaacttatgaaaattga |
| *tetR* | Gene | atgtctagattagataaaagtaaagtgattaacagcgcattagagctgcttaatgaggtcggaatcgaaggtttaacaacccgtaaactcgcccagaagctaggtgtagagcagcctacattgtattggcatgtaaaaaataagcgggctttgctcgacgccttagccattgagatgttagataggcaccatactcacttttgccctttagaaggggaaagctggcaagattttttacgtaataacgctaaaagttttagatgtgctttactaagtcatcgcgatggagcaaaagtacatttaggtacacggcctacagaaaaacagtatgaaactctcgaaaatcaattagcctttttatgccaacaaggtttttcactagagaatgcattatatgcactcagcgctgtggggcattttactttaggttgcgtattggaagatcaagagcatcaagtcgctaaagaagaaagggaaacacctactactgatagtatgccgccattattacgacaagctatcgaattatttgatcaccaaggtgcagagccagccttcttattcggccttgaattgatcatatgcggattagaaaaacaacttaaatgtgaaagtgggtcttaa |
| *mCherry* (*L. lactis*) | Gene | atggtttctaaaggtgaagaagataatatggctatcatcaaagaatttatgcgtttcaaagttcacatggaaggttctgttaatggtcacgaatttgaaatcgaaggtgaaggtgaaggtcgtccttacgaaggtactcaaactgctaaacttaaagttactaaaggtggtcctcttcctttcgcttgggatatcctttctcctcaattcatgtacggttctaaagcctacgttaaacaccctgctgatatccctgattaccttaaactttctttccctgaaggtttcaaatgggaacgtgttatgaatttcgaagatggtggtgttgttactgttactcaagattcttctcttcaagatggtgaatttatctacaaagttaaacttcgtggtactaatttcccttctgatggtcctgttatgcaaaaaaaaactatgggttgggaagcatcttctgaacgtatgtaccctgaagatggtgctcttaaaggtgaaatcaaacaacgtcttaaacttaaagatggtggtcactacgatgctgaagttaaaactacttacaaagctaaaaaacctgttcaacttcctggtgcttacaatgttaatatcaaacttgatatcacttctcacaatgaagattacactatcgttgaacaatacgaacgtgctgaaggtcgtcactctactggtggtatggatgaactttacaaa |
| rrnB T1 | Terminator | ggcatcaaataaaacgaaaggctcagtcgaaagactgggcctttcgttttatctgttgtttgtcggtgaacgctctcctgagtaggacaaatccgccgccctaga |
| rrnB T2 | Terminator | tagctaagcagaaggccatcctgacggatggcctttttgcgtttc |
| **Parts for *E. coli*[S]** | | |
| $P_{CymRC}$ | Promoter | aacaaacagacaatctggtctgtttgtattatggaaaatttttctgtataatagattcaacaaacagacaatctggtctgtttgtattat |
| $P_{J109}$ | Promoter | tttacagctagctcagtcctagggactgtgctagctactagag |

| | | |
|---|---|---|
| $P_{arsOC2}$ | Promoter | cctctgttgacacacattcgttaagtcatatatgtttttgacttatccgcttcgaagaattc |
| $P_{proD}$ | Promoter | cacagctaacaccacgtcgtccctatctgctgccctaggtctatgagtggttgctggataactttacgggcatgcataaggctcgtat aatatattcagggagtccacaacggtttccctctacaaataattttgtttaacttt |
| $P_{oxyS}$ | Promoter | tatccatcctccatcgccacgatagttcatggcgataggtagaatagcaatgaacgattatccctatcaagcattctgactgataatt gctcaca |
| $P_{J23101}$ | Promoter | tttacagctagctcagtcctaggtattatgctagc |
| SarJ | Insulator | agactgtcgccggatgtgtatccgacctgacgatggcccaaaagggccgaaacagtcctctacaaataattttgtttaa |
| RiboJ00 | Insulator | agctgtcaccggatgtgctttccggtctgatgagtccgtgaggacgaaacagcctctacaaataattttgtttaa |
| $RBS_{Ec1}$ | RBS | caccactacccAAGGAGCaacttaa |
| $RBS_{Ec2}$ | RBS | caccactacccTGGGAGCaacttaa |
| $RBS_{Ec3}$ | RBS | caccactacccAGGGAGTaacttaa |
| $RBS_{Ec4}$ | RBS | caccactacccGTGGCGAaacttaa |
| RBS30 | RBS | attaaagaggagaaatactag |
| $sRBS_{arsR1}$ | RBS | aaatacgaccgctttagagaaggcccaca |
| $sRBS_{arsR2}$ | RBS | tactttaactgccagaggccataaaca |
| $sRBS_{arsR3}$ | RBS | ccgctcaatacccgtagacgaata |
| $sRBS_{oxyR}$ | RBS | tgataagtcatctttcgaccttca |
| RBS34 | RBS | ctagtgaaagaggagaaatacta |
| *menD* (*E. coli* BL21) | Gene | atgtcagtaagcgcatttaaccgacgctgggcggcggtcattctggaagcattaacgcgtcacggcgtcagacacatctgtatcg ccccaggctcgcgttctacaccgttaacgttagcggcggcggagaattccgcattcattcaccacacccatttcgatgagcgtggg ttggggcatctggcgctggggctggcgaaagtcagcaagcagccggtggcggtgattgtgacctccggcacggcggtggcaa atctctatccggcactgattgaagccgggttaaccggagaaaaactgattctcttaaccgccgatcgcccgccggagctaattga ctgcggcgcgaatcaggcaattcgccagccgggaatgttcgcctctcaccccacgcacagtatttcattgccgcgcccgaccca ggatatccccgcacgttggctggtttctaccatcgaccacgctctcggtacgcttcatgcgggggggagtccatatcaactgcccgttt gctgaaccgctgtatggcgaaatggacgataccgggcttagctggcaacagcgtctgggtgactggtggcaggacgacaaac cgtggctgcgtgaagcgcctcgtctggaaagtgaaaaacagcgcgactggttcttctggcgacaaaagcgcggcgtggtggttg ccgggcgcatgagtgcggaagagggcaaaaaagttgccctgtgggcgcaaactcttggctggccgctgattggcgatgtgctgt cacaaaccgggcagccgctgccgtgtgccgatctttggttaggcaatgccaaagcgaccagcgagctgcagcaggcgcaaat tgtggtgcaactgggaagcagcctgacggggaaacggctcctgcaatggcaggcaagctgtgaaccagaagagtactggatt gttgatgacattgaagggcgacttgatccggcacaccatcgcggacgtcgcttaattgccaatattgccgactggctggagctgc atccggcagaaaaacgccagccctggtgcgttgaaatcccgcgcctggcggaacaggcaatgcaggcggttattgcccgccg tgatgcgtttggcgaagcgcaactggcgcatcgcatctgcgactatctgcctgaacaggggcaattgtttgttggtaacagcctggt ggtacgtctgattgatgcgctttcgcaacttccggcaggttacccggtgtacagcaaccgtggggccagcggtatcgacgggctg ctttcgaccgccgccggcgttcagcgggcaagcggcaaaccgacgctggcgattgtgggcgatctctccgcactttacgatctca acgcgctggcgttattgcgtcaggtttctgcgccgctggtattaattgtggtgaacaacaacggcgggcaaattttctcgctgttgcc aacgccgcaaagcgagcgtgagcgtttctatctgatgccgcaaaacgtccattttgagcacgccgccgcgatgttcgagctgaa atatcatcgtccgcaaaactggcaggaacttgaaacggcatttgccgacgcctggcgcacgccaaccaccacggtgattgaaa tggtggttaacgacaccgatggtgcgcaaacgctccagcaacttctggcgcaggtaagccatttatga |
| *arsR* | Gene | atgtcatttctgttacccatccaattgttcaaaattcttgctgatgaaacccgtctgggcatcgttttactgctcagcgaactgggagag ttatgcgtctgcgatctctgcactgctctcgaccagtcgcagcccaagatctcccgccacctggcattgctgcgtgaaagcgggct attgctggaccgcaagcaaggtaagtgggttcattaccgcttatcaccgcatattccagcatgggcggcgaaaattattgatgagg cctggcgatgtgaacaggaaaaggttcaggcgattgtccgcaacctggctcgacaaaactgttccggggacagtaagaacattt gcagttaa |
| *oxyR* | Gene | atgaacatccgggacttggaatacttagtagcactcgcggagcaccgtcatttccggagagctgcagatagctgccacgtctccc agccgaccttgagcggacaaattagaaaactcgaagacgaattaggggtgatgttgttagagcgtacttccagaaaagttttgttt acgcaagcgggtatgttgttggtcgatcaagctcgcactgtcttacgtgaagtgaaggtgttgaaggagatggcaagccaacaa ggcgagacaatgagtggccctttacacattggtttgatccccacagtcgggccgtacctcttaccccacatcattccaatgttacac cagacctttcctaaattggagatgtacctccacgaggcgcaaacccaccaattgctcgcgcagctggatagcggcaaattggac tgcgtaatcctcgctctcgttaaggagtccgaggcatttatcgaggtgccgttgttcgatgaacctatgctcttagctatttacgaaga ccacccttgggcgaatcgcgagtgcgttcctatggcagatctggccggggagaaattactgatgttagaggatggccactgcctg cgggatcaggcaatggggttctgctttgaagcgggagcagacgaggatactcattttcgggcgaccagcttagaaacgttgaga |

| | | |
|---|---|---|
| | | aacatggtggcggcaggaagcggaatcactttgttgcccgcattggcggtccctcccgaaagaaaacgcgatggcgtcgtctat ctcccctgtatcaaaccggaacctcgcagaacaatcgggctcgtgtatcgtcctgggagccctttaagatctcggtacgagcagtt ggcagaggcgattcgcgctagaatggacgggcatttcgacaaggtattaaaacaggccgtctaa |
| *mCherry2* (*E. coli)* | Gene | atggtgagtaaaggagaagaaaacaacttagctatcattaaagagttcatgcgcttcaaagttcacatggagggttctgttaacg gtcacgagttcgagatcgaaggcgaaggcgagggccgtccgtatgaaggcacccagaccgccaaactgaaagtgactaaa ggcggcccgctgccttttgcgtgggacatcctgagcccgcaatttatgtacggttctaaagcgtatgttaaacacccagcggatatc ccggactatctgaagctgtcttttccggaaggtttcaactgggaacgcgtaatgaattttgaagatggtggtgtcgtgaccgtcactc aggactcctccctTcaggatggcgagttcatctataaagttaaactgcgtggtactaattttccatctgatggcccggtgatgcagtg taggacgatgggttgggaggcgtctaccgaacgcatgtatccggaagatggtgcgctgaaaggcgaaattaaacagcgcctg aaactgaaagatggcggccattatgacgctgaagtgaaaaccacgtacaaagccaagaaacctgtgcagctgcctggcgcgt acaatgtggatattaaactggacatcttatctcataatgaagattatacgatcgtagagcaatatgagcgcgcggagggtcgtcatt ctaccggtggcatggatgaactatacaaataa |
| L3S2P55 | Terminator | ctcggtaccaaagacgaacaataagacgctgaaaagcgtcttttttcgttttggtcc |
| DT54 | Double terminator | ggaaacacagaaaaaagcccgcacctgacagtgcgggcttttttttttcgaccaaaggctcggtaccaaattccagaaaagaca cccgaaagggtgtttttttcgttttggtcc |
| **Parts for *L. plantarum*[R]** | | |
| $P_{tuf}$ | Promoter + RBS | gcgtgtgtaataagaatttactaacaaaaattcaatttttttgaataatatctgtttacaaatcagattaggctatatataatatttaaggat tctcagtgatgggtgcgcgatttggccttttcactaggatgtagtataatactaactaaagaattgttgagaccattttggcctcgacgt tattcttgcgaaaatcacaggaggtttcatta |
| *sfgfp* (*L. plantarum*) | Gene | atgagcaaaggtgaagaactgtttaccggcgttgtgccgattctggtggaactggatggcgatgtgaacggtcacaaattcagcg tgcgtggtgaaggtgaaggcgatgccacgattggcaaactgacgctgaaatttatctgcaccaccggcaaactgccggtgccgt ggccgacgctggtgaccaccctgacctatggcgttcagtgttttagtcgctatccggatcacatgaaacgtcacgatttctttaaatct gcaatgccggaaggctatgtgcaggaacgtacgattagctttaaagatgatggcaaatataaaacgcgcgccgttgtgaaatttg aaggcgataccctggtgaaccgcattgaactgaaaggcacggattttaaagaagatggcaatatcctgggccataaactggaa tacaactttaatagccataatgtttatattacggcggataaacagaaaaatggcatcaaagcgaattttaccgttcgccataacgtt gaagatggcagtgtgcagctggcagatcattatcagcagaataccccgattggtgatggtccggtgctgctgccggataatcatta tctgagcacgcagaccgttctgtctaaagatccgaacgaaaaacgggaccacatggttctgcacgaatatgtgaatgcggcag gtattacgtaa |

**Supplementary Table 6 | Chemically defined medium (CDM)**

| Component | Final concentration (g/L) |
|---|---|
| **Buffer and salts** | |
| MOPS (3-(N-morpholino)propanesulfonic acid) | 8.371 |
| $K_2HPO_4$ | 0.871 |
| $NH_4Cl$ | 1.070 |
| $Na_2SO_4$ | 1.420 |
| **Carbon source** | |
| Mannitol/glucose | 10 |
| **Metals** | |
| $MgCl_2•6H_2O$ | 0.203 |
| $MnCl_2•4H_2O$ | 0.01 |
| $FeSO_4•7H_2O$ | 0.014 |
| **Amino acids** | |
| Casamino acids | 1.500 |
| Cysteine•HCl•$H_2O$ | 0.145 |
| Tryptophan | 0.050 |
| **Wolfe's Vitamins** | |
| Pyridoxine HCl | 0.001 |
| Thiamine HCl | 0.0005 |
| Riboflavin | 0.0005 |
| Nicotinic acid | 0.0005 |
| Calcium D-(+)-pantothenate | 0.0005 |
| *p*-Aminobenzoic acid | 0.0005 |
| Thioctic acid (α-Lipoic acid) | 0.0005 |
| Biotin | 0.0002 |
| Folic acid | 0.0002 |
| Vitamin B12 | 0.00001 |
| **Wolfe's Minerals** | |
| Nitrilotriacetic acid (NTA) | 0.15 |
| $MgSO_4•7H_2O$ | 0.3 |
| $MnSO_4•H_2O$ | 0.05 |
| NaCl | 0.1 |
| $FeSO_4•7H_2O$ | 0.01 |
| $CoCl_2•6H_2O$ | 0.01 |
| $CaCl_2$ | 0.01 |
| $ZnSO_4•7H_2O$ | 0.01 |
| $CuSO_4•5H_2O$ | 0.001 |
| $AlK(SO)_4•12H_2O$ | 0.001 |
| $H_3BO_3$ | 0.001 |
| $Na_2MoO_4•2H_2O$ | 0.001 |
| **pH adjusted to 6.5** | |

**Supplementary Table 7 | M9 mannitol minimal medium (minimal mM9)**

| Component | Final concentration (g/L) |
|---|---|
| M9 minimal salts (BD Difco) | 11.28 |
| Mannitol | 10 |
| $MgSO_4 \cdot 7H_2O$ | 0.25 |
| $CaCl_2$ | 0.011 |
| Thiamin | 0.0005 |
| Biotin | 0.0002 |
| Casamino acids | 1.5 |
| Tryptophan | 0.05 |

**Supplementary Table 8 | Mannitol MRS medium (mMRS)**

| Component | Final concentration (g/L) |
|---|---|
| Protease Peptone #3 | 10 |
| Tween 80 | 1 |
| Mannitol | 10 |
| Yeast Extract | 5 |
| Potassium phosphate dibasic | 2 |
| Sodium acetate trihydrate | 8.3 |
| Ammonium citrate tribasic | 2.15 |
| Magnesium sulfate anhydrous | 0.1 |
| Manganese sulfate monohydrate | 0.05 |

**Supplementary Table 9 | BRM3 medium for gut microbiota cultivation**

| Component | Final concentration (g/L) |
|---|---|
| **Base** | |
| Tryptone | 1 |
| Proteose peptone #3 | 2 |
| Yeast extract | 2 |
| Sodium chloride | 0.4 |
| Bovine bile | 0.5 |
| Haemin | 0.005 |
| Magnesium sulfate | 0.01 |
| Calcium chloride | 0.01 |
| Tween 80 | 2 mL |
| **Supplement (pH adjusted to 6.8)** | |
| Arabinogalactan | 0.1 |
| D-Cellobiose | 0.15 |
| Maltose | 0.15 |
| D-glucose (Dextrose) | 0.04 |
| Inulin | 0.2 |
| Sodium Bicarbonate | 2 |
| Vitamin K3 | 0.001 |
| Potassium phoisphate dibasic | 0.04 |
| Potassium phosphate monobasic | 0.04 |

**Supplementary Table 10 | Identifiers of enzymes in the DHNA biosynthesis pathway**

| Enzyme | Function | Interpro | Rhea | NCBIfam |
|---|---|---|---|---|
| MenF | Isochorismate synthase | IPR034681 | RHEA:18985 | TIGR00543 |
| MenD | 2-succinyl-5-enolpyruvyl-6-hydroxy-3-cyclohexene-1-carboxylate synthase | IPR004433 | RHEA:25593 | TIGR00173 |
| MenH | 2-succinyl-6-hydroxy-2,4-cyclohexadiene-1-carboxylate synthase | IPR022485 | RHEA:25597 | TIGR03695 |
| MenC | o-succinylbenzoate synthase | IPR010196 | RHEA:10196 | TIGR01928/TIGR01927 |
| MenE | 2-succinylbenzoate--CoA ligase | IPR010192 | RHEA:17009 | TIGR01923 |
| MenB | 1,4-dihydroxy-2-naphthoyl-CoA synthase | IPR010198 | RHEA:26562 | TIGR01929 |
| MenI | 1,4-dihydroxy-2-naphthoyl-CoA hydrolase | IPR030863 | RHEA:26309 | *TIGR00369 |

*TIGR00369 describes hotdog fold thioesterases and was filtered to retain those specifically annotated as 1,4-dihydroxy-2-naphthoyl-CoA hydrolase/thioesterase

### Supplementary References

1. Li, S., De Groote Tavares, C., Tolar, J. G. & Ajo-Franklin, C. M. Selective bioelectronic sensing of pharmacologically relevant quinones using extracellular electron transfer in Lactiplantibacillus plantarum. *Biosensors and Bioelectronics* **243**, 115762 (2024).
2. Meyer, A. J., Segall-Shapiro, T. H., Glassey, E., Zhang, J. & Voigt, C. A. Escherichia coli "Marionette" strains with 12 highly optimized small-molecule sensors. *Nat Chem Biol* **15**, 196–204 (2019).
3. Leloup, L., Ehrlich, S. D., Zagorec, M. & Morel-Deville, F. Single-crossover integration in the Lactobacillus sake chromosome and insertional inactivation of the ptsI and lacL genes. *Appl Environ Microbiol* **63**, 2117–2123 (1997).
4. Li, Q. *et al.* A modified pCas/pTargetF system for CRISPR-Cas9-assisted genome editing in Escherichia coli. *Acta Biochimica et Biophysica Sinica* **53**, 620–627 (2021).
5. Jiang, Y. *et al.* Multigene Editing in the Escherichia coli Genome via the CRISPR-Cas9 System. *Applied and Environmental Microbiology* **81**, 2506–2514 (2015).
